# Identification and characterization of bacterial glycogen-degrading enzymes in the vaginal microbiome

**DOI:** 10.1101/2021.07.19.452977

**Authors:** Dominick J. Jenkins, Benjamin M. Woolston, M. Indriati Hood-Pishchany, Paula Pelayo, Alyssa N. Konopaski, M. Quinn Peters, Michael T. France, Jacques Ravel, Caroline M. Mitchell, Seth Rakoff-Nahoum, Christopher Whidbey, Emily P. Balskus

## Abstract

The healthy human vaginal microbiota is generally dominated by lactobacilli, and the transition to a more diverse community of anaerobic microbes is associated with health risks. Glycogen released by lysed epithelial cells is believed to be an important nutrient source in this environment. However, the mechanism by which vaginal bacteria metabolize glycogen is unclear, with evidence implicating both microbial and human enzymes. Here, we biochemically characterize six glycogen-degrading enzymes (GDEs) from vaginal bacteria that support the growth of amylase-deficient *L. crispatus* on glycogen. We reveal variations in the pH tolerance and susceptibility to inhibition between enzymes from different organisms. Analysis of vaginal microbiome datasets show these enzymes are expressed in all Community State Types. Finally, we confirm the presence and activity of bacterial GDEs in cervicovaginal fluid. This work establishes that bacterial GDEs can participate in the breakdown of glycogen, providing insight into metabolism that may shape the vaginal microbiota.

---

Dysbiosis within the human vaginal microbiota is associated with adverse health outcomes.^1^ The bacterial composition of this community can be classified taxonomically into one of five Community State Types (CSTs).^2^ CST I-III and V are dominated by a single species of *Lactobacillus: L. crispatus, L. gasseri, L. iners*, or *L. jensenii*, respectively. By contrast, CST IV consists of a diverse group of anaerobic and facultative anaerobic microbes, including species of *Gardnerella, Prevotella, Mobiluncus*, and low levels of *Lactobacillus*. The *Lactobacillus*-dominated CSTs are associated with a vaginal pH below 4.5, low Nugent scores, and low levels of inflammation^3^, whereas CST IV is often associated with a higher pH and a number of health sequelae, including HIV acquisition^4^, bacterial vaginosis^5^, and preterm birth.^6^ Thus, the prevailing view of the “healthy” vaginal microbiota is one containing a high proportion of *Lactobacillus* species. However, it is important to note that CST IV is overrepresented in healthy Hispanic and Black women, and is not necessarily indicative of dysbiosis.^7^ It has also recently been revealed that even CSTs dominated by a single species have significant intra-species genetic variation^8^, with multiple strains of the same species co-occurring. The factors contributing to the stability of different vaginal communities, and what causes transitions between the different CSTs, remain poorly understood.^9^ Overall, it has become clear that vaginal microbiota composition alone is insufficient to predict health outcomes, and resolving these questions requires understanding specific functions encoded by vaginal bacteria.

One function known to influence the composition and stability of host-associated microbial communities is the liberation of carbohydrates from specific dietary or host-derived sources by glycoside hydrolase enzymes. This has been well established within the human gut microbiota,^10^ where the presence of an extracellular glycoside hydrolase in *B. ovatus* was beneficial for the success of the organism *in vivo*. ^11^ In addition, co-occurring bacteria have been shown to rely on a glycoside hydrolase ‘producer’ species for community access to carbon sources.^11,12,13^ Compared to the gut, microbial carbohydrate metabolism in the vaginal environment is poorly understood. It is widely believed that glycogen released by exfoliated and lysed vaginal epithelial cells supports the colonization of vaginal lactobacilli^14,15^ since free glycogen levels in vaginal samples have been associated with *Lactobacillus* dominance and a low vaginal pH.^16^ However, until recently, attempts to obtain vaginal *Lactobacillus* isolates that were capable of growth on glycogen were largely unsuccessful.^17,18^ This difficulty raised the important question of how vaginal bacteria access this carbon source.

Glycogen consists of linear chains of α(1-4) linked glucose units, with periodic α(1-6) branches. Metabolism of glycogen requires an extracellular glycoside hydrolase to release shorter glucose polymers (maltodextrins) for import into the cell. Several vaginal lactobacilli have been shown to utilize maltodextrins for growth, leading to the initial hypothesis that a *non-Lactobacillus* glycoside hydrolase in the vaginal environment releases these oligosaccharides.^19^ The detection of human α-amylase via ELISA in cervicovaginal lavage samples (CVLs) by Spear et al. lent support to this proposal, and in the same study those authors demonstrated that amylase-containing human saliva could support *L. crispatus* growth on glycogen.^19^ How human amylase, which is produced predominantly in the pancreas and salivary glands,^19^ comes to be found in genital fluid has not yet been established. Further analysis of the CVLs in that study revealed that most samples had reduced amylase activity at low pH, consistent with the pH profile of human amylase.^20^ In a small fraction of samples, though, activity increased as the pH became more acidic, suggesting the presence of other glycogen-degrading enzymes in the vaginal enviroment.^20^

In addition to human amylase, recent evidence has suggested the presence of other enzymes capable of degrading glycogen in vaginal fluid. The vaginal parasite *Trichomonas vaginalis* secretes multiple glucosidases capable of degrading extracellular glycogen.^21^,^22^ Although this enzyme can support the growth of lactobacilli otherwise unable to grow on glycogen,^21^ this parasite is only found in 2.1% of clinical samples.^23^ Others have demonstrated a potential role for bacterial enzymes in the breakdown of vaginal glycogen. The characterization of a small number of CVLs via proteomics and in-gel amylase assay after native polyacrylamide gel electrophoresis (PAGE) revealed the presence of several putative bacterial glycoside hydrolases, as well as the human enzyme.^24^ In addition, Van der Veer and co-workers isolated several *L. crispatus* strains that could grow on glycogen. These authors suggested a putative secreted Type 1 pullulanase (PulA, EEU28204.2) as the source of amylase activity, based on strain-to-strain variation in its predicted signal peptide which correlates with growth on glycogen.^25,26^

Type 1 pullulanases hydrolyze the α(1-6) linkages in pullulan and other branched oligosaccharides, and this activity enables the release of maltodextrins from highly branched glycogen.^27^ Homologs of PulA are encoded by genomes of various vaginal bacteria,^8^ suggesting this enzyme might not be limited to *Lactobacillus* isolates and highlighting the potential for competition for glycogen and/or cross-feeding. Notably, the proteomics study mentioned above identified putative pullulanases from *Lactobacillus iners* and *Gardnerella vaginalis* in CVLs.^24^ However, the predicted α(1-6) specificity of Type I pullulanases raises questions regarding the fate of the remaining glycogen backbone and how longer branches are hydrolyzed. The identification of these bacterial enzymes also raises questions about the relative role of the human amylase in the vaginal ecosystem. Interpretation of these results is further confounded by the fact that when we began our work, the activities of these reported bacterial enzymes had not been verified biochemically. Clearly, our understanding of glycogen metabolism within the vaginal microbiota would benefit from a detailed characterization of these enzymatic activities.

Here, we report the biochemical characterization of six signal peptide-containing PulA homologs from vaginal microbes representing *Lactobacillus-dominated* CSTs (I and III) and the diverse CST-IV community. We show that these proteins are glycogen-degrading enzymes (GDEs) that support growth of amylase-deficient *L. crispatus* on glycogen. We further characterize the substrate scope of each enzyme, finding that several should be reannotated as amylopullulanases (type II pullulanases). We determine that the *Lactobacillus* GDEs are active at pH 4.0 and have a unique glycogen breakdown product profile compared to other enzymes tested. We demonstrate selective *in vitro* inhibition of GDEs from two vaginal microbes associated with dysbiosis suggesting the possibility of therapeutic interventions for targeted vaginal microbial community modulation. Through bioinformatic analyses of multi-omics datasets, we reveal that the genes encoding these bacterial GDEs are present and transcribed in all CSTs. Finally, using activity-based protein profiling (ABPP),^28^ and an enzymatic assay selective for pullulanase activity – an activity not exhibited by human amylase – we demonstrate that both human and microbial GDEs are present and active in cervical vaginal fluid (CVF). Overall, this work provides molecular insight into the bacterial metabolism of an abundant carbon source in the vaginal microbiota.

## Results

### Identification and purification of bacterial extracellular GDEs

To identify candidate vaginal microbial GDEs, we conducted a BLASTp search of genomes from 151 vaginal isolates in the IMG database (Supplementary File 1) using the putative Type I pullulanase identified by Van der Veer et al in *L. crispatus* (PulA) as our query sequence^25^ (EEU28204.2), with a cut-off of 35% amino acid identity. Hits were further narrowed to those containing a putative signal peptide, since glycogen degradation occurs extracellularly^29^, as well as those with a glycoside hydrolase domain. 62 potential homologs were identified in strains from 11 unique bacterial species (Supplementary File 1), including among others *Lactobacillus crispatus* (7 of 9 strains in the database, average 99% amino acid identity to our query), *Lactobacillus iners* (12 of 13 strains, average 45% identity), *Mobiluncus mulieris* (2 of 4 strains, average 43% identity), *Prevotella bivia* (2 of 2 strains, 40% identity), and *Gardnerella vaginalis* (15 of 18 strains, average 37% identity). Interestingly, gene neighborhood analysis revealed another signal peptide-containing glycoside hydrolase (GH 13) encoded immediately next to the *P. bivia pulA* (25% identity to PulA), so this sequence was included in our study. Each of these bacteria has been previously associated with health or disease, thus subsequent efforts focused on this set of proteins. It should be noted that we also detected potential homologs with lower amino acid identity (Supplementary File 1), including one from *Streptococcus agalacticae* (SAG0089_06185, 33% identity), and one significantly smaller protein in *Gardnerella vaginalis* (HMPREF0424_1317) homologous to a recently reported α-glucosidase enzyme from *Gardnerella* spp. that is active on maltose and other oligosaccharides, but lacks the ability to degrade glycogen.^30^

PFAM domain analysis revealed that all six candidate enzymes contain either an S-layer protein A domain (SlpA), a cell wall binding domain (CWB), or transmembrane helices (TM), suggesting they are located on the cell surface.^31,32,33^ Additionally, each protein contains at least one α-amylase catalytic domain (PF00128), which is a member of the glycoside hydrolase 13 enzyme family known to catalyze the cleavage of various glycosidic bonds.^34^ Interestingly, the *G. vaginalis* enzyme contains two unique amylase domains. In addition, several of the proteins possess putative carbohydrate-binding domains, including the pullulanase domain (PUD; PF03714), which is common among bacterial pullulanase enzymes.^35^ Additional carbohydrate binding modules from the CAZy database found in these enzymes include CBM25 and CBM48, which are responsible for binding different linear and cyclic α-glucans related to starch and glycogen^34,36^ and for multivalent binding to soluble amylopectin and pullulan^37^ (**Fig. 1a**).

**Fig. 1:**
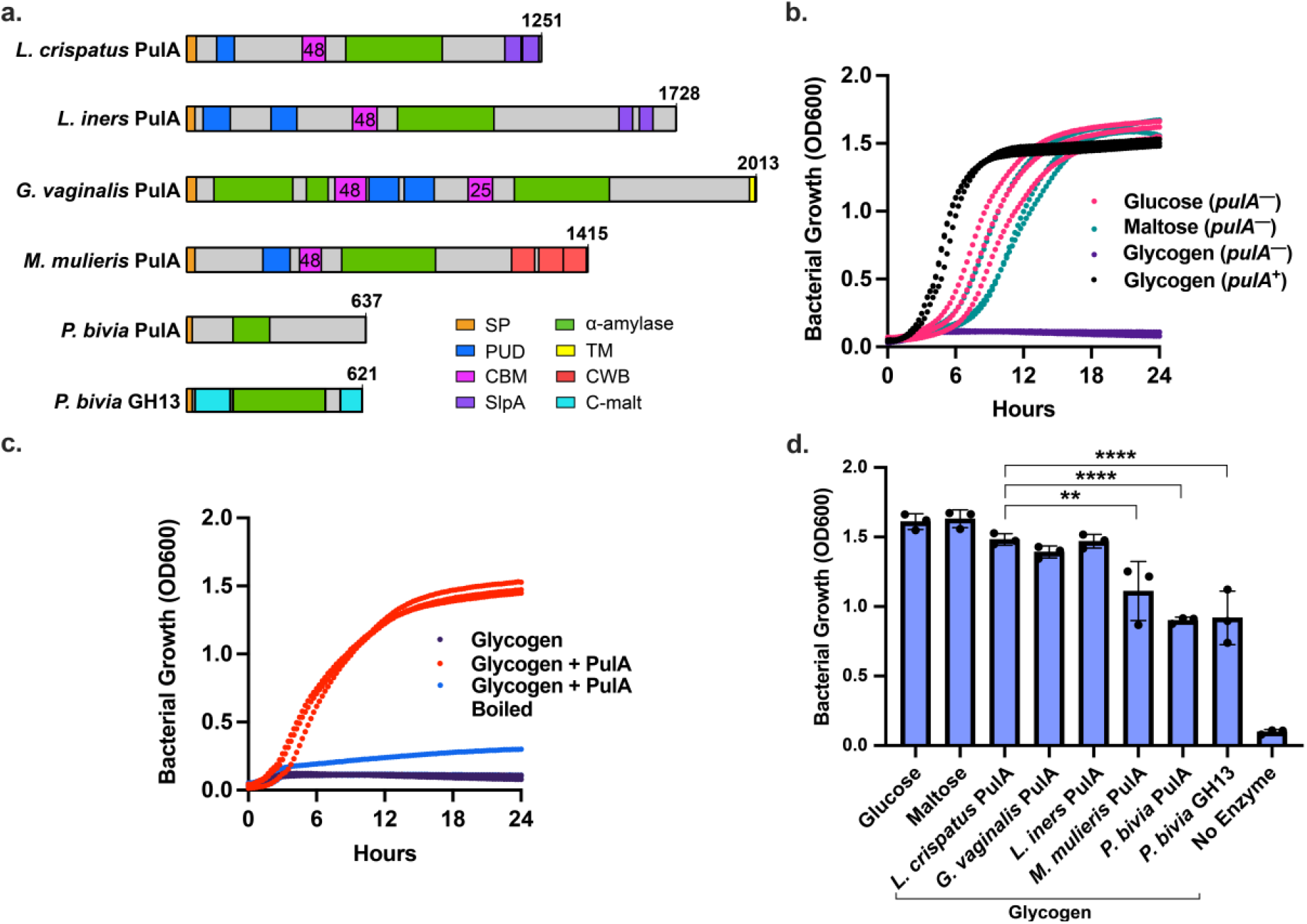
Purified bacterial glycogen-degrading enzymes support *L. crispatus* growth on glycogen. **a.** Predicted domains in putative vaginal microbial extracellular GDEs. Abbreviations: **SP,** Signal peptide; **PUD,** Bacterial pullulanase-associated domain; **CBM,** Carbohydrate binding module; **SlpA,** Surface layer protein A; **α-Amylase**, α-Amylase catalytic domain (GH13); **C-Malt,** Cyclomaltodextrinase domain; **CWB,** Cell wall binding repeat 2; **TM,** Transmembrane helices. **b.** Growth of *L. crispatus* C0176A1 (*pulA*^−^, JAEDCG000000000) and MV-1A-US (*pulA*^+^) on different carbon sources. **c.** *L. crispatus* C0176A1 (*pulA*^−^) grown on oyster glycogen supplemented with 200 nM purified *L. crispatus* PulA. **d.** 24 hr OD600 values of *L. crispatus* C0176A1 (*pulA*^−^) grown on glucose, maltose, or glycogen supplemented with 200 to 400 nM purified protein (***L. crispatus* PulA: *M. mulieris* PulA**, P = 0.0084; ***L. crispatus* PulA: *P. bivia* PulA**, P = <0.0001; ***L. crispatus* PulA: *P. bivia* GH13**, P = <0.0001). All growth curves are representative of three experimental replicates with 3 or 4 technical replicates each day. Error bars represent one standard deviation above and below the mean of all data collected. A multiple comparisons (Tukey) one-way ANOVA was performed to determine statistical significance. P-value symbols: P>0.5 (ns), P≤0.05 (*), P≤0.01 (**), P≤0.001 (***), P≤0.0001(****)

### Purified GDEs support the growth of L. crispatus on glycogen

The domain analysis of these enzymes suggested they act on extracellular carbohydrate polymers. To test this hypothesis, we heterologously expressed and purified each protein, removing the signal peptides for better expression in *E. coli* (**SI Fig. 1, SI Fig. 2**). We then investigated each enzyme’s ability to support growth of a strain of *L. crispatus* that lacks *pulA* in its genome and is unable grow on glycogen (**Fig. 1b; SI Fig. 3**). When purified *L. crispatus* PulA was added to the medium at the time of inoculation, growth on glycogen was recovered (**Fig. 1c; SI Fig. 3**). This provides direct evidence that PulA is sufficient for *L. crispatus* growth on glycogen, as suggested in previous work.^25^ We next expanded this analysis to include the other five purified extracellular glycoside hydrolases. Each enzyme supported the growth of *pulA*-deficient *L. crispatus* on glycogen (**Fig. 1d; SI Fig. 3**), although the optical densities at 600 nm (OD_600_) were significantly lower for *M. mulierus* PulA and both *P. bivia* enzymes compared to the *L. crispatus* PulA, suggesting they are not as efficient at glycogen degradation or that the oligosaccharide products of these enzymes are not as accessible to *L. crispatus* metabolism. Combining the two *P. bivia* enzymes did not affect the overall growth rate (data not shown).

### Kinetic characterization reveals different glycosidic linkage and substrate preferences

Having demonstrated the glycogen degrading activity of these enzymes indirectly through growth complementation, we next performed kinetic characterization using a variety of glucose polymers in order to determine the specificity of each enzyme for the different glycosidic linkages in glycogen and their substrate preference (**Table 1, SI Fig. 4**). In addition to glycogen, we tested amylose, which consists solely of α(1-4) linkages, and pullulan, a polymer that consists of maltotriose units connected by α(1-6) linkages. Consistent with the results of the growth supplementation experiment, all enzymes were active on glycogen. In addition, all were active on pullulan, suggesting the ability to cleave α-1,6 linkages, which was later confirmed by liquid chromatography-mass spectrometry (LC-MS) analysis of the products (discussed below). However, the enzymes differed in their activity toward the α(1-4) linkages in amylose, with *L. crispatus, L. iners, G. vaginalis* and *P. bivia* GH13 enzymes showing activity while the *M. mulieris* PulA and *P. bivia* PulA were inactive (**Table 1, SI Fig. 4**).

**Table 1:** Kinetic analysis of vaginal microbial glycogen-degrading enzymes on various carbohydrate polymers at pH 5.5. Values are representative of three experimental replicates over two days. Error range represents one standard deviation.

| Enzyme | Substrate | $k_{cat}$ ($s^{-1}$) | $K_m$ ($mg\ mL^{-1}$) | Specificity Constant ($mL\ mg^{-1}\ s^{-1}$) | Classification |
| --- | --- | --- | --- | --- | --- |
| *L. crispatus* PulA | Glycogen | $65 \pm 4$ | $0.091 \pm 0.026$ | $710 \pm 210$ | Type II Pullulanase (Amylopullulanase) |
| | Amylose | $33 \pm 7$ | $0.25 \pm 0.14$ | $130 \pm 80$ | |
| | Pullulan | $100 \pm 10$ | $0.15 \pm 0.05$ | $700 \pm 250$ | |
| *L. iners* PulA | Glycogen | $51 \pm 5$ | $0.10 \pm 0.05$ | $500 \pm 220$ | Type II Pullulanase (Amylopullulanase) |
| | Amylose | $30 \pm 4$ | $0.20 \pm 0.07$ | $150 \pm 60$ | |
| | Pullulan | $27 \pm 9$ | $0.93 \pm 0.56$ | $29 \pm 20$ | |
| *G. vaginalis* PulA | Glycogen | $450 \pm 40$ | $0.098 \pm 0.044$ | $4600 \pm 2100$ | Type II Pullulanase (Amylopullulanase) |
| | Amylose | $110 \pm 10$ | $0.27 \pm 0.06$ | $390 \pm 90$ | |
| | Pullulan | $220 \pm 40$ | $0.42 \pm 0.170$ | $520 \pm 230$ | |
| *M. mulieris* PulA | Glycogen | $57 \pm 9$ | $6.1 \pm 1.7$ | $9.5 \pm 3.0$ | Type I Pullulanase |
|  | Amylose | NA | NA | NA |  |
| | Pullulan | $150 \pm 20$ | $0.33 \pm 0.14$ | $440 \pm 200$ | |
| *P. bivia* PulA | Glycogen | $0.81 \pm 0.31$ | $11 \pm 7$ | $0.077 \pm 0.058$ | Type I Pullulanase |
|  | Amylose | NA | NA | NA |  |
| | Pullulan | $60 \pm 6$ | $0.24 \pm 0.07$ | $250 \pm 70$ | |
| *P. bivia* GH13 | Glycogen | $5.1 \pm 0.60$ | $3.8 \pm 1.0$ | $1.4 \pm 0.3$ | Type II Pullulanase (Amylopullulanase) |
| | Amylose | $210 \pm 50$ | $0.68 \pm 0.33$ | $310 \pm 160$ | |
| | Pullulan | $120 \pm 30$ | $1.5 \pm 0.6$ | $79 \pm 38$ | |

In general, the measured kinetic parameters were consistent with prior values observed for bacterial enzymes that process these substrates (glycogen^38,39,40^, amylose^41,42^, pullulan^43,41^). Comparing the specificity constants (*k*_cat_/K_m_) for each substrate revealed that glycogen is the preferred substrate for the *G. vaginalis* and *L. iners* PulA enzymes. The *L. crispatus* PulA had similar specificity constants for both pullulan and glycogen, with a lower specificity for amylose. Other enzymes, including *P. bivia* PulA and *M. mulierus* PulA, had higher specificity constants for pullulan in comparison to glycogen and amylose, while the *P. bivia* GH 13 enzyme preferred amylose (**Table 1; SI Fig. 4**). Taken together, these data demonstrate that both *Lactobacillus* PulA enzymes, the *G. vaginalis* PulA enzyme, and the glycoside hydrolase from *P. bivia* can cleave α-1,4 linkages and likely also α-1,6 linkages needed for the complete utilization of glycogen, and support their reassignment as type II pullulanases or amylopullulanases (EC. 3.2. 1.1/41, reviewed in^44^). In contrast, the lack of activity of the *M. mulieris* and *P. bivia* PulA enzymes on amylose identifies them as type I pullulanases and may explain their reduced ability to complement *L. crispatus* growth on glycogen (**Fig. 1d**).Table 1: Kintetic analysis of vaginal microbial glycogen degrading enzymes

### *Site-directed mutagenesis of* G. vaginalis *PulA confirms the contribution of both active sites to enzyme activity*

*G. vaginalis* PulA contains two α-amylase catalytic domains, suggesting there may be two functional active sites. To probe the activity of each domain individually, active site mutants of each catalytic domain as well as a double active site mutant were constructed by mutating the critical catalytic aspartate residue to alanine (D233A, D1317A, **SI Fig. 5**). The specific activities of the single active site mutants were significantly reduced, retaining approximately 5% of wild-type activity, and the double mutant was completely inactive (**SI Fig. 5**). This result confirms the individual activity of each active site and suggests they may act synergistically to process glycogen.

### *The* Lactobacillus *GDEs maintain activity at low pH*

*Lactobacillus-dominant* communities are typically associated with a lower vaginal pH than mixed anaerobic communities due to their high production of lactic acid, which excludes other microbes.^16^ We therefore hypothesized that GDEs from lactobacilli may have evolved to maintain activity at a lower pH than those from other vaginal bacteria. Measuring specific activity on glycogen, we screened a pH range from 2.5 to 8.0 (**Fig. 2**). Five of the GDEs exhibited maximum activity at a pH between 5.5 and 6.0, which is consistent with values observed for other characterized bacterial pullulanases and amylopullulanases.^45^ *P. bivia* PulA exhibited maximum activity at a slightly lower pH between 4.5 and 5 (**Fig. 2**). We observed that most of the enzymes from vaginal anaerobes – *G. vaginalis* PulA, *M. mulieris* PulA, and *P. bivia* GH 13 – have almost no activity at pH 4.0, which is within the range of a healthy vaginal community.^46^ Critically, however, the *Lactobacillus crispatus, Lactobacillus iners*, and *Prevotella bivia* PulAs display 34%, 51% and 97% of their maximal activity at pH 4.0, suggesting these enzymes are better suited for a low pH environment compared to enzymes from other species tested. The ability to liberate oligosaccharides from host-derived glycogen at low pH may explain how vaginal lactobacilli (*L. iners and L. crispatus*) can acquire a key carbon source under these conditions, potentially contributing to their dominance.

**Fig. 2:**
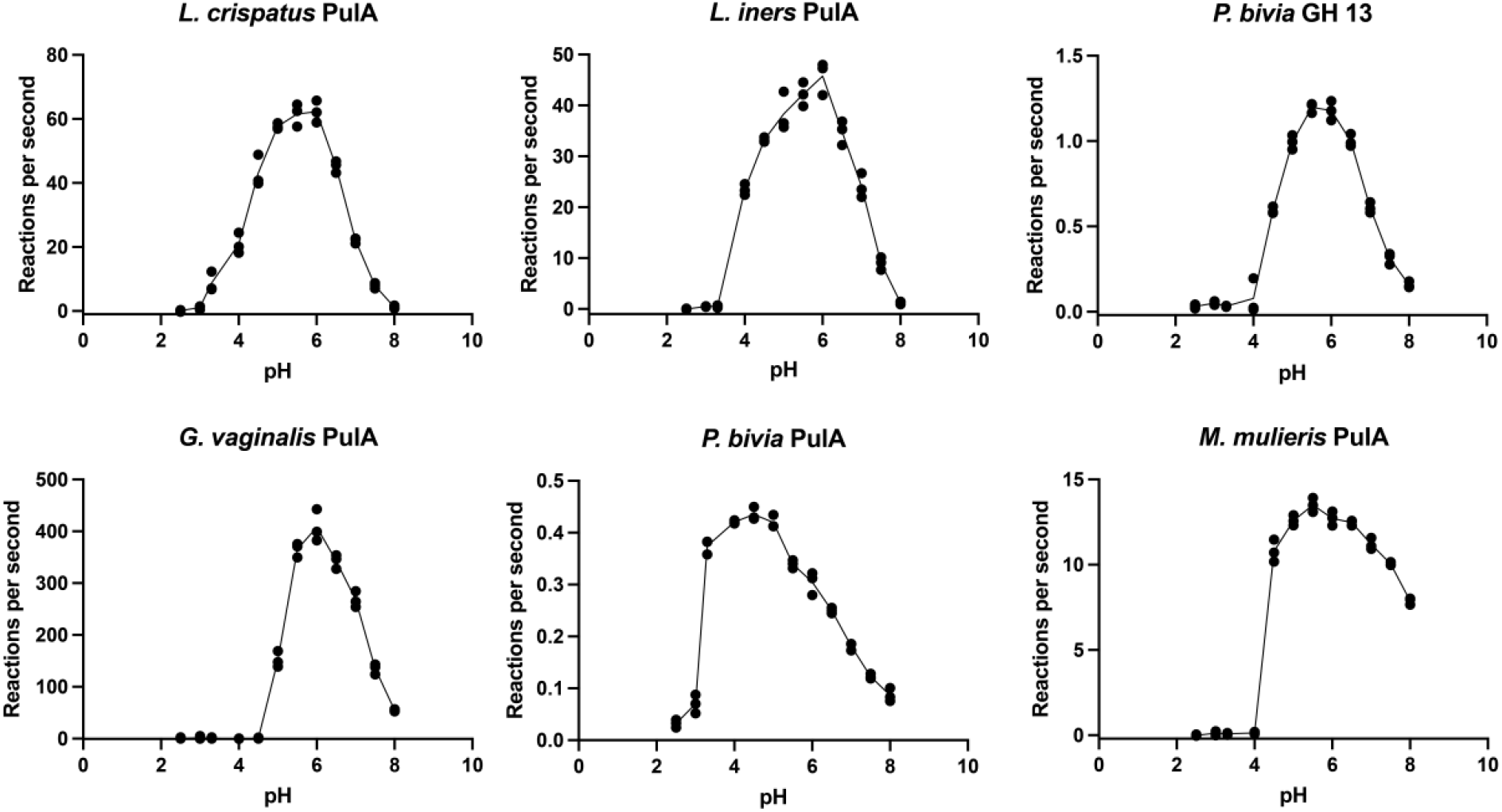
*Lactobacillus* amylopullulanases are adapted to a low pH environment. pH profiles of six extracellular glycogen-degrading enzymes. Buffer systems consisted of glycine (pH = 2.5 to 3.3), sodium acetate (pH = 4.0 to 5.5), MES (pH = 6.0 to 6.5), and HEPES (pH = 7.0 to 8.0). Data is representative of three experimental replicates over two days.

### LC-MS analysis reveals distinct oligosaccharide breakdown product profiles by GDEs

Next, we sought to characterize the oligosaccharides produced by each enzyme. Following overnight incubations with glycogen, amylose, and pullulan, oligosaccharide production was quantified by LC-MS (**Fig. 3**). Both human amylase and the *P. bivia* GH 13 produced predominantly maltose and a relatively small amount of glucose from glycogen and amylose. In contrast, the enzymes annotated as Type I Pullulanases (PulA homologs) produced longer oligosaccharides in addition to maltose, including maltotriose and in some cases maltotetraose. These results are similar to those observed for previously characterized bacterial amylopullulanses.^47,44^ Amongst the pullulanases, maltotetraose was not detected in the *G. vaginalis* PulA reaction, and was only detected at a low level in *M. mulieris* and *P. bivia*. However, the *Lactobacillus-derived* PulA enzymes produced a higher relative amount of maltotetraose when acting on amylose or glycogen. During incubation with pullulan, all of the bacterial enzymes produced predominantly maltotriose, however the human salivary amylase was not active. The sole production of maltotriose is common among enzymes that degrade pullulan^43,45,47^ and provides direct confirmation that vaginal bacterial GDEs can cleave α-1,6 linkages. This observation further corroborates the functional assignments of these enzymes as either amylopullulanases or pullulanases. Notably, this experiment verified biochemically, that pullulanase activity is unique to bacterial GDEs in the vaginal environment and is not exhibited by the human enzyme (**Table 1, Fig. 3**).

**Fig. 3:**
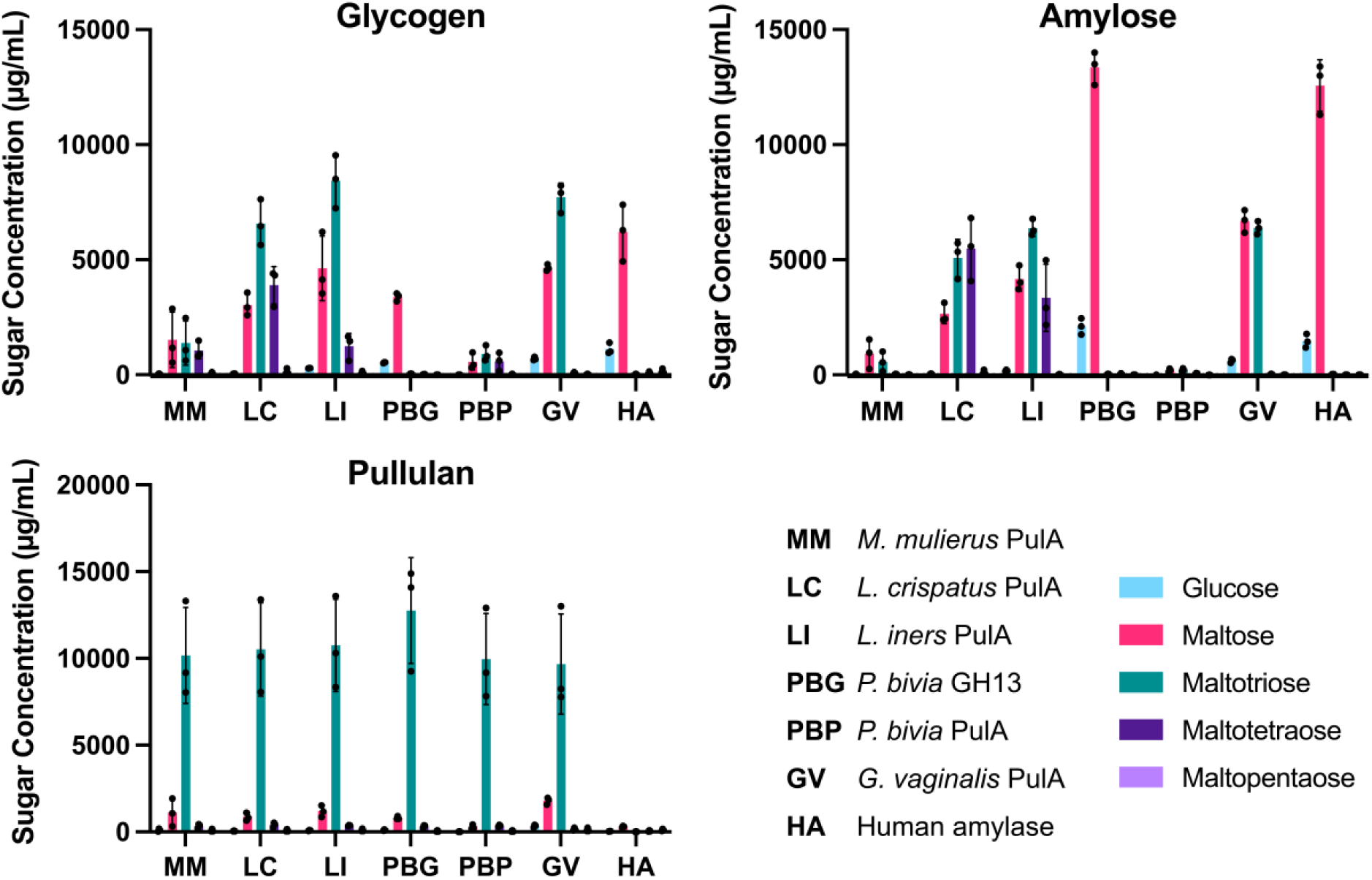
GDEs from different vaginal lactobacilli produce unique breakdown product profiles. Polymer breakdown products generated following overnight incubation with purified enzyme. LC-MS analysis is representative of three experimental replicates performed over three days. Error bars represent one standard deviation above and below the mean. Protein abbreviations are as follows, **MM**, *M. mulierus* PulA; **LC**, *L. crispatus* PulA; **LI**, *L. iners* PulA; **PBG**, *P. bivia* GH13, **PBP**, *P. bivia* PulA; **GV,** *G. vaginalis* PulA; **HA**, Human Amylase

### Acarbose selectively inhibits GDEs from G. vaginalis and P. bivia

Given their role in enabling growth on glycogen, we hypothesized that these enzymes may be good targets for possible therapeutic intervention aimed at establishing a *Lactobacillus*-dominant community. The variability between these microbial GDEs in their pH profiles, glycogen breakdown product profiles, and kinetic parameters, led us to propose that they may have enough structural and functional variation to be selectively inhibited. We therefore attempted to identify inhibitors for the GDEs from microbes associated with dysbiosis by screening purified enzymes against a panel of four clinically relevant human amylase inhibitors. Of the compounds tested, only acarbose and acarviosin showed any inhibition (**Fig. 4a**). We then determined IC50 values for the inhibition of each enzyme by acarbose, the most promising inhibitor. Acarbose inhibited *G. vaginalis* PulA, *P. bivia* PulA, and *P. bivia* GH 13 enzymes with IC50 values of 120 ± 30 μM, 420 ± 90 μM, and 0.84 ± 0.05 μM, respectively, while the *L. crispatus, L. iners*, and *M. mulierus* enzymes were largely unaffected (**Fig. 4b**). For comparison, the IC_50_ of acarbose towards human amylase is approximately 11 μM.^48^

**Fig. 4:**
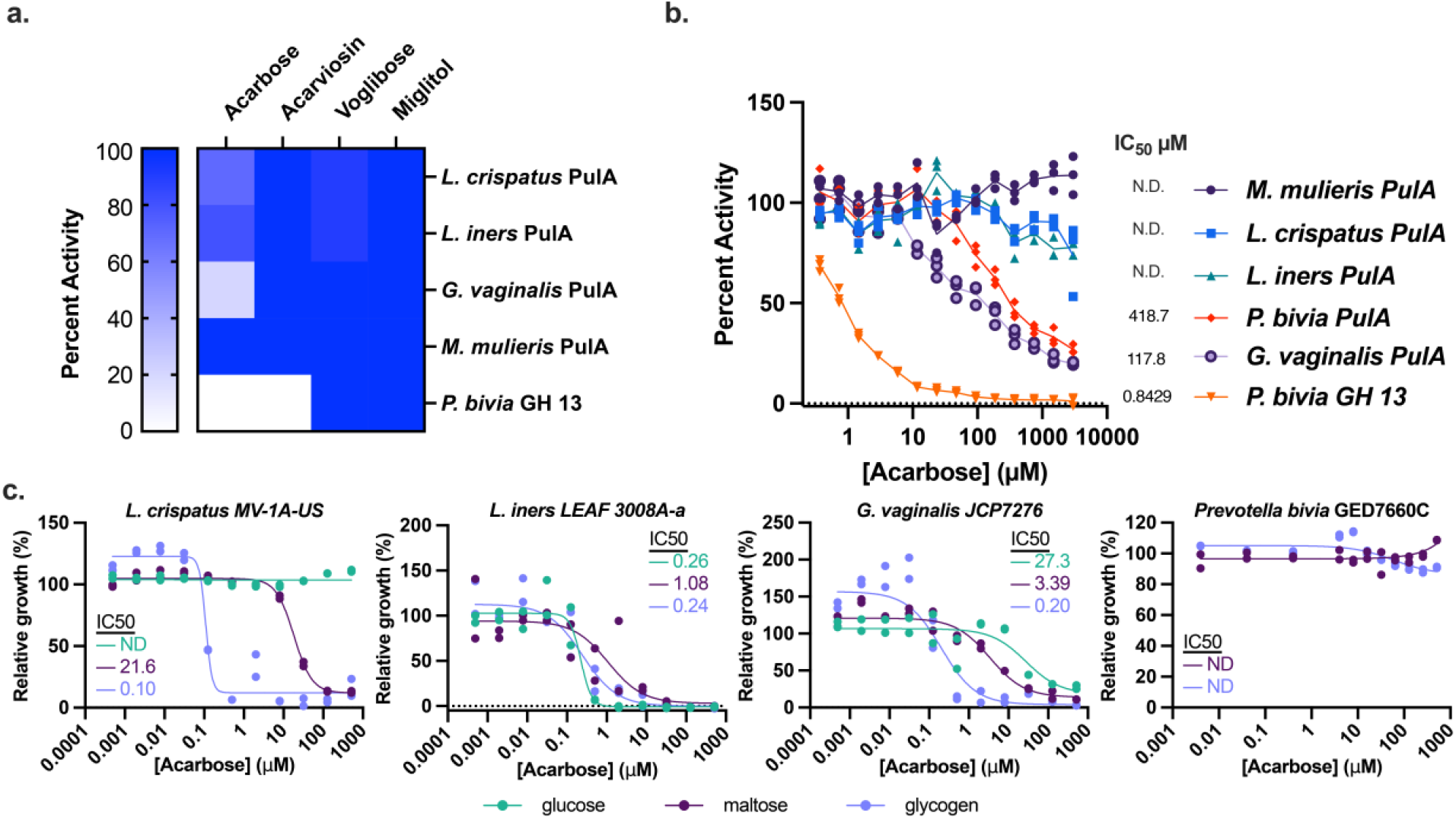
Selective inhibiton of vaginal microbial GDEs. **a.** Effects of known amylase inhibitors (1 mM) on the activities of purified GDEs toward a BODIPY fluorescent starch substrate (n=1). **b.** Inhibitory activity of acarbose toward purified extracellular amylases. A BODIPY fluorescent starch substrate was used and activity was normalized to a no inhibitor control. Data are representative of three experimental replicates over two days. **c.** Bacteria were grown in the presence of the indicated concentrations of acarbose in media containing either glucose, maltose or glycogen as the primary carbohydrate source. Growth in the presence of inhibitor was normalized to the untreated control. IC_50_ values were calculated using a least-squares regression of the normalized values. ND (Not determined) is indicated when the resulting curve fit was poor and an IC_50_ value could not be confidently determined, or the overall growth inhibition was less than 10%. Data are representative of at least two biological replicates performed over two days.

Since acarbose selectively inhibited GDEs from CST IV-associated microbes, we characterized its effect on the growth of several vaginal microbes as a first step toward testing its utility for community modulation. While acarbose inhibited *G. vaginalis* growth on glycogen (IC_50_ = 0.2 μM), it also inhibited *L. crispatus* growth on maltose (IC_50_ = 22 μM) and glycogen (IC_50_ = 0.1 μM) even though *L. crispatus* PulA was not affected *in vitro*. Interestingly, *L. crispatus* growth was not affected when glucose was the primary carbon source (**Fig. 4c**). These data suggest there are additional *L. crispatus* enzymes involved in maltodextrin metabolism that are inhibited by acarbose. Despite potently inhibiting the *P. bivia* GH13 *in vitro*, acarbose had no impact on *P. bivia* growth on any substrate. (**Fig 4c**). This suggests that *P. bivia* PulA, which was less inhibited *in vitro*, is likely the predominant source of extracellular amylase activity in this organism, and that, unlike in *L. crispatus*, intracellular maltodextrin catabolism in *P. bivia* is not affected by acarbose. Overall, although acarbose is not a suitable candidate for community modulation due to its broad target spectrum, these results highlight differences between the glycogen-degrading enzymes that may be targeted for selective inhibition in future efforts.

### Bacterial GDEs are present in human vaginal microbiome datasets

Having identified *bona fide* GDEs in genomes of sequenced vaginal bacterial isolates, we next sought to understand if these genes are present and expressed in the vaginal environment. While other searches have detected putative microbial amylases in clinical samples using proteomics,^24^ metagenomics, and metatranscriptomic searches,^49,26^ we employed shortBRED (Short, Better Representative Extract Dataset) to identify biochemically characterized GDEs in a previously generated database of 178 paired vaginal metagenome and metatranscriptomes from 40 non-pregnant, reproductive age women who self-collected vaginal swabs over 10 weeks.^49,50^ ShortBRED is a computational tool that identifies unique amino acid sequences that are distinct to a query protein (85% identity cutoff) and uses these sequences to quantify reads that encode the protein in sequence datasets.

Combined reads encoding our six biochemically characterized GDEs were more abundant in CST I metagenomes compared to CST II and CST IV metagenomes (**Fig. 5a**). This increased abundance in CST I samples is due to reads encoding *L. crispatus pulA* which is consistent with the taxonomic classification of this CST (**Fig. 5b**). *L. crispatus pulA* was detected in 84.6% of the metagenomes and 89.7% of the metatrascriptomes from participants classified as CST I. *M. mulieris pulA* was not detected in these individuals and the other four genes encoding GDEs were detected in less than 11% of both metagenomes and metatranscriptomes (**SI Figure 6**). In CST III, which is dominated by *L. iners, L. iners pulA* was detected in 41.9% and 32.3% of the metagenomes and metatranscriptomes, respectively. Interestingly, genes encoding other GDEs (*L. crispatus* PulA, *G. vaginalis* PulA, both *P. bivia* GDEs) were detected in greater than 20% of the metagenomes classified as CST III. However, the detection of these genes in the metatranscriptomes was highly variable (6.45 %-38.7%). This indicates the GDEs from multiple species are frequently present in this CST.

**Fig. 5:**
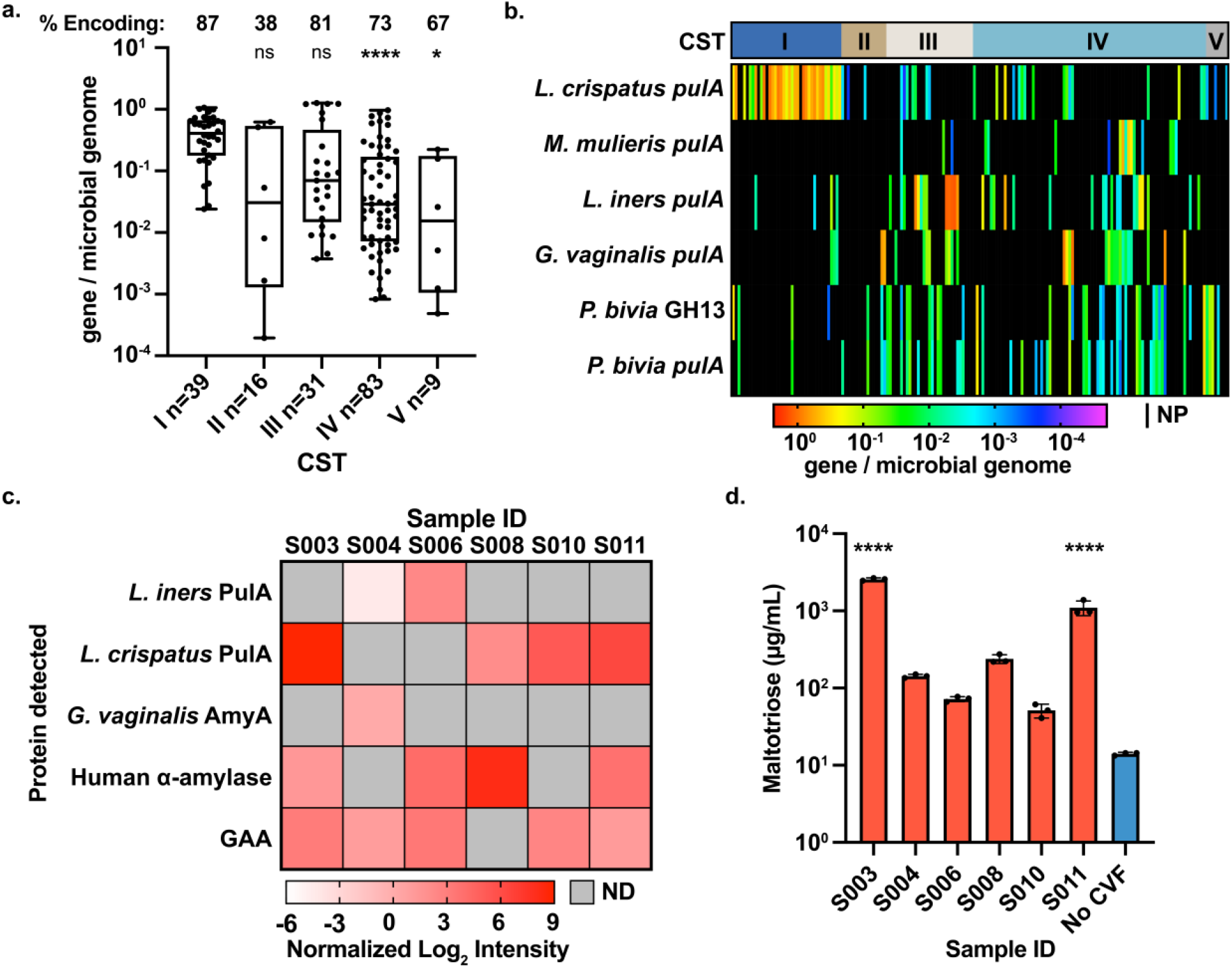
Human CVFs samples contain active human amylase and microbial GDEs. **a.** Metagenomic analysis of 168 participant samples using ShortBRED analysis of biochemically characterized GDEs stratified by CST. Only samples encoding a microbial GDE were plotted. **% Encoding,** represents the percentage of samples that contain > 0 genes / microbial genome. A multiple comparisons (Dunnett) one-way ANOVA was performed to determine statistically significant differences compared to CST I abundance. (CST IV, P = < 0.0001; CST V, P = < 0.0196) The box represents 1.5 of the interquartile range and the whiskers represent the minimum to the maximum of the dataset. **b**. Heat map of metagenomic presence and abundance detected using ShortBRED within each sample. **C.**ABPP analysis identifies microbial GDEs and human proteins (α-amylase and GAA) in CVF supernatants. Abbreviations: **ND**, Not detected **GAA**, Lysosomal α-glucosidase **d.** Human CVF contains distinctly microbial pullulanase activity at pH 5.5. Data are representative of three experimental replicates over two days and the error bars are one standard deviation above and below the mean. A multiple comparisons (Dunnett) one-way ANOVA was performed to determine statistically significant differences compared to the no CVF sample (Blue) (S003, P = <0.0001; S011, P = <0.0001). P-value symbols: P>0.5 (ns), P≤0.05 (*), P≤0.01 (**), P≤0.001 (***), P≤0.0001(****)

All GDEs were detected in CST IV samples, at frequencies ranging from 16.9% to 36.1% of sample metagenomes. While *L. crispatus pulA* was detected in 39.8% of CST IV metatranscriptomes, the sequences of GDEs from other species were detected in only 3.6% to 10.8% of the datasets (**SI Figure 6**). Overall, this analysis confirms that the enzymes we identified are present in vaginal metagenomes and expressed in a broad range of vaginal microbial CSTs, with CST III and CST IV communities in particular harboring GDEs from multiple species.

### Analysis of clinical samples reveals activity of human and microbial enzymes

Having characterized microbial GDEs that are expressed in human vaginal microbiomes (**Fig. 5ab**), we next sought to detect the activity of these enzymes and the human amylase in human clinical samples. We initially analyzed 20 CVL sample supernatants spanning a range of Nugent scores (0-8). First, we compared total amylase activity in the samples to the concentration of human amylase, as determined via ELISA (**SI Fig 7, SI Fig. 8**). We assayed for amylase activity toward a starch substrate across a range of pHs (4.4, 5.5, and 6.8). pH 4.4 was chosen because it is characteristic of a healthy *Lactobacillus* dominated vaginal environment, pH 5.5 corresponds to the maximal activity of most microbial enzymes we tested (**Fig. 2**), and pH 6.8 is the pH optimum of the human amylase. At all pH values examined, there was a statistically significant correlation between these measurements (**SI Fig. 8)** suggesting that the majority of the amylase activity in these samples is of human origin. However, it is interesting to note that at pH values below the pH optimum of the human enzyme, the correlation coefficient is reduced (pH 4.4, r = 0.8556, 95% CI 0.6472 to 0.9450; pH 5.5, r = 0.9611, 95% CI 0.8966 to 0.9857; pH 6.8, r = 0.9713, 95% CI 0.9252 to 0.9891) suggesting other enzymes could be contributing to this activity at low pH (**SI Fig. 8).** When comparing these results to the Nugent scores (Low Nugent (0-3); High Nugent (7-10)), we found no difference in amylase activity or human amylase levels between these two groups (**SI Fig. 9**).

To rule out the possibility that the human amylase ELISA signal arose from cross-reactivity of the antibodies with conserved structural features of one or more of the microbial GDEs identified here, we assayed our purified enzymes using the same kit and found no cross-reactivity at enzyme concentrations as high as 1 μM (**SI Fig. 7**). Though we cannot rule out the possibility that the antibodies react with additional microbial amylases not identified here, these data strongly corroborate previous findings of human amylase in vaginal samples and suggest that the contribution of this host enzyme in shaping the vaginal microbiota should not be overlooked, despite the existance of microbial enzymes with related activities.

We next attempted to determine if any of the microbial GDEs were present in human samples by assaying for activity directly in a different group of CVF samples using ABPP.^51^ We utilized fluorescent (Cy 5)- and biotin-tagged activity-based probes designed to label inverting α-glucosidases and amylases (**SI Fig. 10**).^24,52^ For the α-amylase probes (Amy-ABP) the reporter group is located on the C4’ position of a maltose analog while α-glucosidase probes (Glc-ABP) have the reporter group attached to the aziridine nitrogen of a glucose analog, mimicking attachment to the C1 position (**SI Fig. 10**).^28,52^ To ensure that our proteins of interest reacted with these probes, we incubated the six purified GDEs with the Cy5 labeled probes and detected reactivity using fluorescent SDS-PAGE analysis. All the enzymes characterized in this study reacted with the Amy-ABP probe *in vitro*, and every enzyme, with the exception of *L. crispatus* and *M. mulierus* PulA reacted with the Glc-ABP probe (**SI Fig. 11**).

After confirming the reactivity of our probes with *bona fide* microbial GDEs, CVF supernatants were labeled with biotin-tagged probes followed by a pull-down, tryptic digest, and LC-MS/MS analysis of the peptides. Three proteins enriched by the Amy-ABP probe were of bacterial origin and were derived from *L. crispatus* PulA, *L. iners* PulA, and *G. vaginalis* AmyA (**Fig. 5c, SI Fig. 12**). *L. crispatus* PulA and *L. iners* PulA were mutually exclusive, consistent with previous studies finding their cooccurrence uncommon.^7^ In one sample (S004), we did observe co-occurrence of two microbial enzymes: *L. iners* PulA and *G. vaginalis* AmyA. Although we did not characterize *G. vaginalis* AmyA in this study, it is interesting to note that this enzyme contains a signal peptide suggesting it most likely metabolizes extracellular polysaccharides (**SI Fig. 12**). In a recent preprint, the catalytic domain from a homolog of *G. vaginalis* AmyA was characterized *in vitro* (77% ID full length protein, 99% ID for the catalytic domain).^53^ Intriguingly, we also detected human α-amylase (AMY1) in the majority of samples (**Fig. 5c, SI Fig. 12**). No pattern regarding microbial-microbial or microbial-human enzyme co-occurrence is apparent, which may be due to the small number of samples tested.

In contrast to the Amy-ABP-enriched proteins, all of the Glc-ABP-enriched proteins were human in origin. The protein with the highest intensity signal was lysosomal α-glucosidase, which is canonically localized to the lysosome, but has been detected in CVF in other studies (**Fig. 5c, SI Fig. 12, Supplementary File 2**).^54–56^ Overall, these data demonstrate that multiple bacterial GDEs are present and active in the vaginal environment including *L. crispatus* PulA, *L. iners* PulA and *G. vaginalis* AmyA.

To further validate the activity of microbial GDEs in these samples, we used a LC-MS based assay. This assay leverages the observation that all GDEs we characterized metabolize pullulan, whereas the human amylase does not (**Fig. 3**). Every CVF sample we tested showed a time-dependent enzymatic production of maltotriose from pullulan (**SI Fig. 13**). Notably, the two most active samples, S003 and S011, had the highest intensity of *L. crispatus* PulA in the ABPP experiment (**Fig. 5c**) and generated significantly increased levels of maltotriose compared to a no CVF control (**Fig. 5d**). These results further demonstrate that vaginal microbial GDEs contribute to carbohydrate breakdown activity in clinical samples and highlight the utility of an accessible assay for microbial GDEs that does not depend on proteomic workflows.

## Discussion

In this study, we biochemically characterized six glycogen-degrading enzymes from vaginal bacteria. Our results demonstrate that, in addition to relying on human amylase, some vaginal bacteria possess alternative enzymes for accessing glycogen. These findings are further validated by a separate report of the glycogen-degrading activity of *L. crispatus* PulA (GlgU, 99% ID to PulA).^57^ A critical finding of our study examining GDEs from a variety of vaginal bacteria is that, despite sharing a common annotation, the substrate preferences of the different PulAs are quite distinct. While the *L. crispatus, L. iners* and *G. vaginalis* amylopullulanases have the highest specificity for glycogen, enzymes from other organisms are most active on amylose or pullulan and have comparatively high K_m_ values for glycogen. This is consistent with the unique carbohydrate binding modules found in each protein and may suggest adaptation to process structurally distinct glucose polymers in the vaginal environment. Further, since the oligosaccharides produced from glycogen breakdown are released extracellularly and may act as ‘public goods’^58^, the differences in the product distributions of these enzymes may suggest that differential availability of glycogen-derived oligosaccharides between community state types supports the growth of distinct non-glycogen-degrading bacteria via cross-feeding, as has been shown within the gut microbiota^13^. A better understanding of the structure of glycogen within the vaginal environment, and whether it differs among CSTs, is needed to further evaluate this possibility.

Our work also suggests a potential mechanism supporting *L. crispatus* growth and dominance. Specifically, we discovered that the *L. crispatus* amylopullulanase is active at the low pH values (~3.5-4) associated with vaginal health. This enzymatic activity may therefore enable *L. crispatus* to access glycogen under conditions where the human amylase is minimally active and the growth of competing bacteria is inhibited. Critically, the pH profiles of amylases cannot be predicted from primary sequence analysis,^59^ further highlighting the need for biochemical characterization to support bioinformatic interrogations of microbial functions within the human microbiome.

Though our work shows that *Lactobacillus crispatus* encodes an enzyme that allows growth on glycogen, interesting questions remain about how other common vaginal lactobacilli lacking a *pulA* homolog can dominate a community, for example *Lactobacillus jensenii* (CST V) and *Lactobacillus gasseri* (CST II). Possibly these microbes rely on cross-feeding of oligosaccharides produced by human amylase or GDEs from other vaginal microbes, since they are capable of growth on a range of maltooligosaccharides.^19^

Our efforts to identify inhibitors for bacterial GDEs implies that these enzymes have sufficient structural diversity to be selectively targeted. Unfortunately, growth assays in bacterial cultures suggest these existing amylase inhibitors have additional effects on maltodextrin metabolism that limit their ability to target specific bacteria. Future work could examine other differences in maltooligosaccharide catabolism between vaginal strains, and phenotypic screening approaches may help identify inhibitors that favor the growth of *Lactobacillus* strains through the collective inhibition of amylases from both bacteria associated with dysbiosis and the host.

Finally, we characterized GDEs in human vaginal microbiome samples using a combination of bioinformatic analyses and approaches that directly detect active enzymes (ABPP and pullulanase activity assay). Our results show that the genes encoding microbial GDEs are not only found in vaginal microbial genomes, but are also expressed and active in their native environment. This represents a notable milestone in our understanding of this bacterial metabolic activity. Previous bioinformatic analyses did not distinguish between intracellular and extracellular enzymes and aimed to capture all encoded enzymes having a predicted glycogen-debranching activity.^49^ In contrast, all enzymes used as a query in our analysis are predicted to be extracellular and have been biochemically confirmed to degrade glycogen, increasing confidence that they possess this activity in the vaginal environment.^34^ The higher metagenome abundances of characterized microbial GDEs in CST I samples compared to CST IV samples suggests a potential link between the abundance of these enzymes and health status that warrants further investigation.

A major challenge in detecting microbial amylase activity in clinical samples has stemmed from a lack of ability to distinguish human amylase activity from microbial amylase activity because most analyses employ a substrate that can be utilized by both enzymes.^24^ Here, we developed and validated a simple biochemical assay for pullulanase activity that rapidly identifies microbial-specific enzyme activity in clinical samples without the contribution from the human enzyme.^19,24^ Finally, our results, and those from other recent efforts, ^19,24^ show that the presence of bacterial GDE activity in clinical samples is highly variable, highlighting a need to test larger numbers of better characterized clinical samples. We anticipate our pullulanase activity assay will find broad utility in the analysis of such samples and enable further study of the biological roles of bacterial GDEs.^60^

Overall, the insights gained from this investigation highlight the need for complementing bioinformatic analysis with detailed biochemical characterization of vaginal microbial enzymes. This improved understanding of the activities of vaginal bacterial GDEs will enable future exploration of bacterial glycogen metabolism in the vaginal microbiome and its contribution to community composition, stability, and dysbiosis.

## Acknowledgements

We thank Amelia Woo for help cloning several of the bacterial PulA homologs, as well as Beverly Fu for critical reading of the manuscript. We are grateful to Agnes Bergerat-Thompson at Massachusetts General Hospital for providing CVL samples. Financial support for this study was provided to E.P.B by the Bill & Melinda Gates Foundation under award No. OPP1189211. E.P.B. is a Howard Hughes Medical Institute Investigator. M.I. H.-P. was supported as a Fellow in the Pediatric Scientist Development Program, Award Number HD000850 from the Eunice Kennedy
Shriver National Institute of Child Health and Human Development and through a Physician Scientist Fellowship from the Doris Duke Charitable Foundation (Grant # 2019129). S.R.-N. is supported by a Career Award for Medical Scientists from the Burroughs Wellcome Fund, a Pew Biomedical Scholarship, a Basil O’Connor Starter Scholar Award from the March of Dimes, 1K08AI130392-01, and by the NIGMS/NIH under award DP2GM136652. C.W., A.K.N., and M.Q.P. were supported by the M.J. Murdock Charitable Trust under award NS-201913756 and the Seattle University College of Science and Engineering. Proteomics experiments were supported by the Proteomics & Metabolomics Shared Resource of the Fred Hutch/University of Washington Cancer Consortium (P30 CA05704). P.P. was supported by the National Science Foundation Graduate Research Fellowship (NSF-GRFP). M.T.F. and J.R. were supported by the Bill and Melinda Gates Foundation award No. OPP1189217. We thank Dr. Hermann Overkleeft and Dr. Gideon Davies for their generous gift of the ABPP probes. We thank all members of the Gates Foundation Vaginal Microbiome Research Consortium for the helpful conversations about the work, and David Relman for the gift of the *L. crispatus* C0176A1.

## Author Contributions

E.P.B., B.M.W., S.R.-N., and M.I.H.-P. conceived the study. D.J.J. and B.M.W. designed and conducted enzyme purification and biochemical characterization experiments. E.P.B., D.J.J., B.M.W., C.W., S.R.N., and M.I.H.-P. wrote the manuscript. D.J.J. and M.I.H. designed and conducted bacterial growth experiments. C.W., A.N.K., and M.Q.P. designed and conducted ABPP experiments. D.J.J., P.P., and E.P.B. designed bioinformatic analysis of metagenomic and metatranscriptomic data. P.P. conducted bioinformatic analysis of multiomics sequencing data. M.T.F. and J.R. assisted with access, analysis, and interpretation of the metagenomic and metatranscriptomic data. C.M. provided CVL samples and corresponding metadata. All authors contributed to the interpretation of data, were involved in the revision of the manuscript, and approved the final manuscript. D.J.J. and B.M.W. contributed equally to the study.

## Methods

### Reagents

Unless otherwise noted, commercial chemicals were of the highest purity available and purchased from Sigma-Aldrich. Acetonitrile for LC-MS was purchased from Honeywell-Burdick & Jackson.

### Identification and cloning of glycogen degrading enzymes

Homologs of PulA in *L. crispatus*^25^ (EEU28204.2) were identified by BLASTp searches of genomes from vaginal isolates in the IMG database^61^ using an E-value cut-off of 1×10^-5^. The IMG database contains 151-vaginal isolate genomes from the Human Microbiome Project with a sample body subsite of vaginal (Supplementary File 1). Hits were further curated by removing proteins with no predicted signal peptide (SignalP v5.0^62^). A panel of six candidates with >35% amino acid identity from microbes associated with health or disease were selected for further study. The strains were obtained, and genomic DNA was extracted with a DNeasy UltraClean Microbial Kit (Qiagen). Genes were amplified via PCR with primers designed to remove the signal peptide (**SI Fig. 2**) and cloned into the *E. coli* expression vector pET28a (Novagen) via Gibson assembly (New England Biolabs, NEB) to generate an N-terminal His6-tagged gene. All plasmids were verified via Sanger sequencing (Eton Biosciences) and transformed into the expression host BL21 (DE3) (*P. bivia* enzymes) or ArcticExpress (DE3) (all other enzymes) for expression and purification. Complete lists of plasmids and primers are provided in Supplementary Tables 1 and 2, respectively.

### Purification of glycogen degrading enzymes

Cultures containing expression plasmids were grown in 2-6 L of LB (Research Products International, RPI) containing 50 μg/mL kanamycin. Once cultures reached an optical density at 600 nm (OD_600_) of 0.6-0.8, they were cooled to 15 °C and induced with 250 μM IPTG (Teknova). After 16 h at 15 °C, cells were harvested (6720 x g for 10 min at 4 °C) and the pellets were stored at –20 °C until use. Pellets were resuspended in 98% Buffer A (50 mM HEPES, 300 mM KCl, 10% glycerol, pH 7.8), 2% Buffer B (50 mM HEPES, 300 mM KCl, 10% glycerol, 500 mM imidazole, pH 7.8) supplemented with EDTA-free protease inhibitor cocktail (Sigma). Cells were lysed via homogenization (3 x 15,000 psi, Emulsiflex-C3, Avestin) and lysates were clarified (16,000 x g for 45 min at 4 °C) before being loaded onto a 5 mL HisTrap column (GE Healthcare). This was followed by one column volume (c.v.) of 2% Buffer B, and 2 c.v. of 10% Buffer B. Protein was eluted using a linear gradient from 10 to 100% Buffer B over 20 c.v., and protein-containing fractions and purity was determined by SDS-PAGE (BioRad). The following day, protein-containing fractions were pooled, concentrated to a volume of approximately 1 mL in a spin concentrator (Millipore), and purified by size exclusion chromatography (GE Healthcare, Superdex 200) in 100% buffer A. Fractions were again analyzed by SDS-PAGE and protein-containing fractions were pooled, concentrated (Millipore, Amicon 30 kDa), flash frozen in liquid nitrogen, and stored at –80 °C until use. Protein concentration was determined using a Bradford assay (BioRad). Molecular weights used for concentration determination were calculated in EXPASY using the predicted protein sequences.

### L. crispatus *growth recovery assay with purified protein*

MRS broth containing glucose (BD Difco) was prepared according to the manufacturer’s protocol. For growth assays on different carbon sources, MRS broth without glucose (Food Check Systems, pH 6.5-6.6) was prepared according to the manufacturer’s recipe and supplemented with either 2%D-glucose (Sigma), 2% maltose monohydrate (VWR), or 5% glycogen from oyster (Sigma). Each media type was filter sterilized (0.2 μm) and left inside an anaerobic chamber with an atmosphere of 2.5 % H_2_, 5 % CO_2_, 92.5 % N_2_ (Coy Labs) overnight for equilibration. Starter cultures of *L. crispatus* C0176A1 (*pulA*^−^) and *L. crispatus* MV-1A-US (*pulA*^+^) were inoculated into MRS media (BD Difco) in Hungate tubes (VWR) and incubated overnight at 37 °C. The next day, purified protein was thawed and added to 5% glycogen MRS media to a concentration ranging between 200-400 nM. After protein addition, the media was again filter sterilized before use. As a negative control, protein boiled at 100 °C for 15 min was also used in the assay. Immediately after protein addition, 50 μL of each media type was aliquoted into a 384-well TC-treated, clear microplate (Corning). 1 μL of overnight culture was used to inoculate each well. The plate was sealed (VWR), and growth was monitored in a plate reader (Biotek) inside of an anaerobic chamber (Coy Labs) for 24 h by measuring OD_600_ every 15 min while incubating at 37 °C. Growth conditions contained 2.5% H_2_, 97.5 % N_2_ with oxygen levels maintained below 20 ppm. Three experimental replicates over three days were performed with three to four technical replicates on each day.

### Kinetic analysis of glycogen-degrading enzymes

Kinetic analysis of glycogen-degrading enzymes was performed using a previously described reducing sugar assay,^43^ modified for a 96-well format. 300 μL reactions were set up containing substrate (0.0048-10 mg mL^-1^ glycogen; 0.0012-1.25 mg mL^-1^ Pullulan (Megazyme); or 0.0048-1.25 mg mL^-1^ amylose in a final concentration of 2% DMSO), 0.8-700 nM enzyme, and reaction buffer (20 mM sodium acetate, pH 5.5, 0.5 mM CaCl_2_). Reactions were incubated at 37 °C for 15 min and 50 μl aliquots were removed (2, 5, 7.5, 10, 15 min) into 125 μL of the BCA stop solution (0.4 M sodium carbonate, pH 10.7, 2.5 mM CuSO_2_, 2.5 mM 4,4’-dicarboxy-1,2’-biquinoline, 6 mM L-serine). After 30 min incubation at 80 °C, 125 μL was transferred to a TC-treated flat bottom plate (Greiner Bio) and absorbances were read at 540 nm. A maltose standard curve (0.000610-0.625 mg mL^-1^) was used to quantify hydrolysis activity. Initial velocities were calculated via linear regression selecting the data points that produced the highest initial rate utilizing at least three data points. *K_M_* and *k_cat_* parameters were determined by fitting the Michaelis-Menten equation to the initial velocity data using nonlinear regression (Graphpad Prism 8). Replicates consisted of three experimental trials over two days.

### *Enzyme pH profile and* G. vaginalis *PulA active site mutant activity on Glycogen*

The reducing sugar assay described above was used to determine the dependence of activity on pH. Reactions contained 1.25 mg mL^-1^ glycogen, 0.9 to 850 nM enzyme, and assay buffer ranging in pH from 2.5 to 8.0 (20 mM glycine, 0.5 mM CaCl_2_, pH 2.5-3.3; 20 mM sodium acetate, 0.5 mM CaCl_2_, pH 4.0-5.5; 20 mM MES, 0.5 mM CaCl_2_, pH 6.0-6.5; 20 mM HEPES, 0.5 mM CaCl_2_, pH 7.0-8.0). Initial velocities were determined for each condition and normalized to enzyme concentration. A maltose standard curve (0.000610-0.625 mg mL^-1^) was used to quantify hydrolysis. Replicates consisted of three experimental replicates over two days. *G. vaginalis* PulA active site mutants were constructed using a multi-fragment Gibson assembly amplified from the wild type expression vector pETpullGV using the primers listed in Supplementary Table 2. pETpullGV-AS1 (ΔAS1) and pETpullGV-AS2 (ΔAS2) contained a D233A and D1317A mutation respectively designed to inactivate the catalytic aspartate of the amylase domains of these proteins. pETpullGV-DM (ΔDBL) contained both mutations. Specific activities of *G. vaginalis* PulA active site mutants were determined as described above, at pH 5.5.

### Polysaccharide breakdown product analysis and growth studies

To measure polysaccharide breakdown products, reactions were set up containing 10 mg mL^-1^ substrate and 500 nM enzyme, all dissolved in assay buffer (20 mM sodium acetate, 0.5 mM CaCl_2_, pH 5.5) and incubated at 37 °C overnight. The next day, samples were quenched with a 10-fold dilution into 90% LC-MS grade acetonitrile. The plates were centrifuged (3220 x g for 10 min, 4 °C) and the samples were diluted 1000-fold into LC-MS grade acetonitrile before analysis by UHPLC-MS using a Xevo TQ-S (Waters) operating in MS/MS mode with electrospray ionization (ESI). 5 μL of sample was injected onto an Acquity BEH/Amide UPLC Column (Waters, 1.7 μm, 130Å, 2.1 mm x 50 mm) heated to 40 °C. A flow rate of 0.5 ml min^-1^ was used, with the following gradient: 0-1.0 min at 97 % B (acetonitrile with 0.1% formic acid) and 3% A (H_2_O with 0.1% formic acid) isocratic, 1.0-4.0 min 97-30% B, 4.0-5.0 min at 30% B isocratic, 5.0-5.1 min at 30-97% B, 5.1-7.0 min at 97% B isocratic. Carbohydrate products were detected by ESI in positive mode (capillary voltage 3.10 kV; cone voltage 42 V; source offset voltage 50 V; desolvation temperature 500 °C; desolvation gas flow 1000 L/hr; cone gas flow 150 L/hr; nebuliser 7.0 bar). See supplementary information for compound-specific detection parameters (**SI table 3**). For quantification, standards of glucose, maltose (VWR), maltotriose (Carbosynth), maltotetraose (Carbosynth), and maltopentaose (Carbosynth) were prepared ranging from 0.001-10 μg mL^-1^ in 9:1 acetonitrile:water. Oligosaccharide peak areas were quantified using the standard curve and the data was normalized to a no-enzyme control to account for non-enzymatic substrate breakdown. These data consisted of three experimental replicates run over three different days.

### Growth assays with amylase inhibitors

Inhibition growth assays were performed in an anaerobic chamber (Coy Labs) with an atmosphere of 2.5 % H_2_, 5 % CO_2_, 92.5 % N_2_. Bacteria were inoculated from single colonies into a peptone-yeast extract base broth (PYTs, pH 7.0-7.2) consisting of proteose peptone (20 g L^-1^), yeast extract (10 g L^-1^), MgSO_4_ (0.008 g L^-1^), K_2_HPO_4_ (0.04 g L^-1^), KH_2_PO_4_ (0.04 g L^-1^), NaHCO_3_ (0.4 g L^-1^), vitamin K (0.0025 g L^-1^), hemin (0.005 g L^-1^), L-cysteine • HCl (0.25 g L^-1^), Tween 80 (0.25 mL L^-1^), horse serum (50 mL L^-1^), and glucose (2 g L^-1^), and incubated at 37 °C for approximately 24 h. Cultures were adjusted to OD_600_ 0.4-0.5, sub-cultured at a 1:50 dilution into fresh PYTs (without glucose) with the indicated carbohydrates added to a final concentration of 2 g L^-1^. Glycogen was from oyster (Sigma, G8751). Assays were performed in duplicate in 384-well plates, sealed with BreathEasy gas permeable membranes (Diversified Biotech) under anaerobic conditions. Bacterial growth was monitored by measuring the OD_600_ at 1 h intervals for 48 h in a BioTek Epoch2 plate reader. Data are representative of at least two independent experiments performed on separate days. Data were normalized to blank (uninoculated) media. For inhibition assays, bacteria were cultivated as above, with the addition of acarbose at the indicated concentrations. Half maximal inhibitory concentrations (IC_50_) were calculated using relative growth data determined by measuring the OD_600_ of cultures grown with each tested concentration of inhibitor taken at the time an untreated control reached stationary phase. Data were then normalized to the OD_600_ of the untreated control, and IC_50_ values were calculated using a least squares regression (GraphPad Prism 8).

### Metagenomics and metatranscriptomics

ShortBRED was used to quantify the abundance of the six biochemically characterized microbial GDEs in vaginal metagenomes and metatranscriptomes sequenced by the Vaginal Microbiome Research Consortium (VMRC).^50^ First, ShortBRED-Identify was used to create markers for all 6 PulA sequences using UniRef90 2017 as a reference list (Supplementary File 3), and an 85% cluster ID setting. After markers were generated, they were used in ShortBRED-Quantify to determine the abundance of *pulA* genes in paired metagenome and metatranscriptome databases generated by the VMRC (Bioproject PRJNA797778). The scripts used for processing the datasets have been previous described.^49^ We analyzed 187 paired metagenomes and metatranscriptomes from 40 participants. The output from ShortBRED-Quantify is reads per million reads per kilobase million (RPKM) and this was normalized to counts per microbial genome using the average genome sizes (AGS) of each metagenome sample, calculated using MicrobeCensus.^63^ We normalized the output from ShortBRED using the previously derived equation shown below.^64^

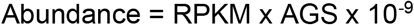

Sample metadata was used to bin the results by community state type (CSTI n=39, CSTII n=16, CSTIII n=31, CSTIV n=83, CSTV n=9). The fraction of samples positive for a microbial GDE gene in a given CST was calculated by dividing the number of samples that contained a hit (Reads > 0) by the total number of samples with the corresponding CST. Plots were generated using Prism 9.4.

### CVL analysis for amylase activity and human amylase abundance

CVLs were obtained from Dr. Caroline Mitchell at Massachusetts General Hospital (IRB: 2014P001066). We obtained informed consent from all donors and complied with all relevant ethical regulations. All donors were white and between the ages of 21 and 63. One donor identified as Hispanic. The donors had a variety of diagnoses and methods of contraception, and Nugent scores ranged from 0 to 8 (**Supplementary Table 4**). CVLs were collected using 3 mL of sterile saline washed over the cervix and vaginal walls with a transfer pipette and then re-aspirated. Samples were centrifuged (10,000 x g for 10 min at 4 °C) and the supernatants were decanted and used in the assay. Purified proteins were diluted in Buffer A (50 mM HEPES, 300 mM KCl, 10% glycerol, pH 7.8) to 1 μM, then used in the assay. Human salivary amylase was purchased from Sigma Aldrich (A1031-1KU). Human amylase was detected in CVLs using an ELISA for human pancreatic amylase (Abcam ab137969) according to manufacturer’s instructions. Two experimental replicates were conducted over two days, and two technical replicates of the standard curve were also measured each day.

Amylase activity of CVL supernatants was determined using the EnzCheck Ultra Amylase Assay Kit (ThermoFisher, E33651). The fluorescent substrate was prepared according to the kit instructions using three different buffers (20 mM sodium acetate, 0.5 mM CaCl_2_, pH 4.4; 20 mM sodium acetate, 0.5 mM CaCl_2_, pH 5.5; 20 mM MES, 0.5 mM CaCl_2_, pH 6.8). 10 μL of CVL supernatant was added to each well of a clear-bottom black 96-well plate and then diluted with 40 μL of pH adjusted buffer. The reactions were initiated with 50 μL of substrate and incubated for 30 min at 37 °C. The pH adjusted buffer made up 90% of the reaction volume and each kit reagent was dissolved in the corresponding buffer. Readings were taken every 51 s by monitoring an excitation of 485 nm and an emission of 528 nm. Initial rates were calculated in the plate reader software (Biotek) by determining the highest slope that covered at least 5 data points.

### Activity-based protein profiling in CVF samples

CVF samples were collected from Seattle University affiliates (IRB: FY2022-002). We obtained informed consent from all donors and complied with all relevant ethical regulations. All six donors were between the ages of 18-25 years old. One donor self-identified as Hispanic, two as multi-racial (Hispanic/white and East Asian/white), and three as white. None were pregnant or menopausal. Three were using hormonal birth control and three were not. Four reported the day of their menstrual cycle, ranging from day 9 to day 21 (**Supplementary Table 5**). Donors self-collected a sample by inserting a Soft Disc and then waiting 1-4 hours before removing the disc and placing it into a 50 mL conical vial. Within one hour of collection, CVF was removed from the disc through the addition of 1 mL of PBS and centrifugation at 200 rpm for eight minutes. Samples were then frozen in 0.1 mL aliquots at –70 °C.

Biotinylated and fluorescent probes for α-amylases (CYR1114 and CYR232 respectively)^28^ and α-glucosidases (JJB384 and JJB383 respectively)^52^ were kindly provided by Dr. Hermann Overkleeft and Dr. Gideon Davies (Leiden University). Prior to use, CVF samples were spun down (10,000 x g for 5 min) to remove mucins. Protein concentration of all CVF supernatants was determined by BCA Assay. CVF samples were normalized to a protein concentration of 1 mg/mL using sterile PBS and EDTA-free protease inhibitor cocktail (Roche) was added. Normalized CVF supernatant was then incubated with the fluorescent amylase probe at a final concentration of 25 μM or the fluorescent glucosidase probe at a final concentration of 10 μM. Negative controls included no probe controls in which vehicle (1% v/v DMSO in water) was added instead of probe and heat shock controls to identify background fluorescence or off-target labeling. Samples were incubated for 4 hours at 37 °C after which 4x Laemmli buffer was added and proteins were denatured by heating at 95 °C for 5 minutes. Proteins were then separated on a 4-20% PAGE gel (Bio-Rad) and probe fluorescence was visualized (Azure C600). Following visualization of the fluorescent probe, gels were stained with Bio-Safe Coomassie stain (Bio-Rad) to visualize total protein content.

Active amylases were enriched using ABPP and identified via LC-MS/MS as previously described, with slight modifications.^51^ CVF supernatant as prepared as above was divided into three 400 μL aliquots. Biotinylated amylase probe (final concentration 25 μM), biotinylated glucosidase probe (final concentration 10 μM), or an equal volume of vehicle (1% DMSO in water) was added, and samples were incubated for 4 hours at 37 °C. After labeling, 400 μL of ice-cold methanol was added and samples were stored at −70 °C overnight to precipitate proteins. Precipitated protein was collected via centrifugation (10,000 x g for 10 min), redissolved in of 500 μL of 1.2% SDS in PBS, and heated at 95 °C for 2 minutes. Samples were then centrifuged (14,000 x g for 5 min) to remove insoluble proteins.

100 μL of streptavidin agarose resin (Thermo Fisher) was prepared by washing with 0.5% w/v SDS in PBS (three times), 6 M urea in 25 mM NH_4_HCO_3_ (three times), and PBS (three times) using a vacuum manifold. Washed resin in 2 mL of PBS was then added to protein samples and samples were incubated rotating at 37 °C for 1 hour. Samples were then transferred to columns (Bio-Rad Poly-Prep) on a vacuum manifold and washed with 1 mL volumes of 0.5% w/v SDS in PBS (three times), 6 M urea in 25 mM NH4HCO3 (three times), ultrapure water (three times), PBS (nine times), and 25 mM NH_4_HCO_3_ (five times). Resin was then transferred in 6 M urea in 25 mM NH_4_HCO_3_ to low-bind Eppendorf tubes and reduced with 5 mM DTT at 37 °C for 30 minutes followed by alkylation with 10 mM iodoacetamide at 50 °C for 1 hour. Samples were washed with PBS (nine times) and 25 mM NH_4_HCO_3_ (five times). Resin was then transferred to new low-bind Eppendorf tubes, resuspended in 200 μL of 25 mM NH_4_HCO_3_, and 0.4 μL of 0.25 μg/μL trypsin (Promega, proteomics grade) in 25 mM HEPES was added. Samples were incubated overnight at 37 °C rotating. Supernatants were collected followed by an additional resin wash with 150μL of 25 mM NH_4_HCO_3_, which was added to the original supernatant. The peptides were then dried down (Speed-vac) prior to further analysis.

Except for S010, liquid chromatography tandem-mass spectrometry (LC-MS/MS) analysis was performed with a Thermo Scientific Easy1200 nLC (Thermo Scientific) coupled to a tribrid Orbitrap Eclipse (Thermo Scientific) mass spectrometer. In-line de-salting was accomplished using a reversed-phase trap column (100 μm × 20 mm) packed with Magic C18AQ (5-μm 200Å resin; Michrom Bioresources) followed by peptide separations on a reversed-phase column (75 μm × 270 mm) packed with ReproSil-Pur C18AQ (3-μm 120Å resin; Dr. Maisch, Baden-Würtemburg, Germany) directly mounted on the electrospray ion source. A 60-minute gradient using a two-mobile-phase system consisting of 0.1% formic acid in water (A) and 80% acetonitrile in 0.1% formic acid in water (B). The chromatographic separation was achieved over a 60 min gradient from 8 to 30% B over 57 min, 30 to 45% B for 10 min, 45 to 60% B for 3 min, 60 to 95% B for 2 min and held at 95%B for 11 min at a flow rate of 300 nL/minute. A spray voltage of 2300 V was applied to the electrospray tip in line with a FAIMS source using varied compensation voltage −40, −60, −80 while the Orbitrap Eclipse instrument was operated in the data-dependent mode, MS survey scans were in the Orbitrap (Normalized AGC target value 300%, resolution 240,000, and max injection time 50 ms) with a 1 sec cycle time and MS/MS spectra acquisition were detected in the linear ion trap (Normalized AGC target value of 50% and injection time 35 ms) using HCD activation with a normalized collision energy of 27%. Selected ions were dynamically excluded for 60 seconds after a repeat count of 1. For S010, peptide samples were disolved in 2% acetonitrile in 0.1% formic acid (20 μL) and analyzed (18 μL) by LC/ESI MS/MS with a Thermo Scientific Easy-nLC 1000 (Thermo Scientific) coupled to a tribrid Orbitrap Fusion (Thermo Scientific) mass spectrometer. In-line de-salting was accomplished using a reversed-phase trap column (100 μm × 20 mm) packed with Magic C18AQ (5-μm 200Å resin; Michrom Bioresources) followed by peptide separations on a reversed-phase column (75 μm × 250 mm) packed with ReproSil-Pur 120 C_18_ AQ (3-μm 120Å resin Dr. Maisch, Germany) directly mounted on the electrospray ion source. A 90-minute gradient from 2% to 35% acetonitrile in 0.1% formic acid at a flow rate of 300 nL/minute was used for chromatographic separations. A spray voltage of 2200 V was applied to the electrospray tip and the Orbitrap Fusion instrument was operated in the data-dependent mode, MS survey scans were in the Orbitrap (AGC target value 500,000, resolution 120,000, and injection time 50 ms) with a 3 sec cycle time and MS/MS spectra acquisition were detected in the linear ion trap (AGC target value of 10,000 and injection time 35 ms) using HCD activation with a normalized collision energy of 27%. Selected ions were dynamically excluded for 20 seconds after a repeat count of 1.

Samples were analyzed using FragPipe IonQuant enabled.^65–68^ Spectra were matched to a database containing UniProt human reference proteins; UniRef90 proteins for *L. crispatus, L. iners, L. gasseri, L. jensenii, G. vaginalis, A. vaginae, P. bivia*, and *M. mueleris;* common contaminants; and reverse protein sequences as decoys for FDR estimation (accessed 25 May 2022). Raw data are available in Supplementary File 4. Abundance data were analyzed using Perseus.^69^ Abundance data were log2 transformed and normalized using width adjustment. For S010, protein groups present in two of three replicates were averaged, and the data tables were combined. Proteins with at least a 2-fold increased abundance relative to the No Probe control in one biological sample, 2 spectral counts across all samples, and a ProteinProphet probability of above 0.95 (corresponding to an approximately 2% FDR) were searched for CAZyme domains using dbCAN 2.^70^

### Pullulanase activity assays in CVF samples

5 μL of CVF fluid (not centrifuged) was added to 95 μL of 10 mg mL^-1^ pullulan (Megazyme) in assay buffer (20 mM sodium acetate, 0.5 mM CaCl_2_, pH 5.5). The reaction mixtures were incubated at 37 °C and timepoints at 3, 5, 8, and 24 h were taken by diluting 100-fold into 9:1 acetonitrile:water. Samples were further diluted 1,000-fold in LC-MS grade acetonitrile and analyzed by LC-MS as described above for the detection of maltotriose. Samples were normalized to a no enzyme control. Three experimental replicates over two days were performed for each sample.

### Inhibitor screening and IC50 determination for acarbose

The inhibitory effect of a panel of four small-molecule inhibitors was determined using a modification of the amylase activity assay in CVLs used above (EnzCheck Ultra Amylase Assay Kit ThermoFisher, E33651). For initial screening, enzymes were preincubated with 1 mM acarbose (Abcam), acarviosin (Toronto Research Products), voglibose (Spectrum Chemical), or miglitol (Tokyo Chemical Industry) for 15 min at room temperature. For IC_50_ analysis, enzyme (2.5-50 nM) was preincubated for 15 min at room temperature with acarbose (Abcam) ranging from 0.366 μM to 3000 μM in a total volume of 50 μL. The reactions were initiated with 50 μL of substrate and incubated for 30 min at 37 °C monitoring with an excitation of 485 nm and an emission of 528 nm. Initial rates were calculated in the plate reader software (Biotek) by determining the highest slope that covered at least 8 data points. Percent activity was calculated by normalizing the activity to a no inhibitor control. IC_50_ values were calculated using a nonlinear fitting of the data to the inhibitor vs normalized response function (GraphPad Prism 8). Error associated with the IC_50_ values represents 95% confidence intervals.

### Data Availability

The protein identification number for each enzyme characterized is as followed, *L. crispatus* PulA (EEU28204.2), *L. iners* PulA (EFQ51965.1), *G. vaginalis* PulA (EPI56559.1), *M. mulieris* PulA (EEZ90738.1), *P. bivia* PulA (WP_061450340.1), *P. bivia* GH 13 (WP_036862728.1). The *L. crispatus* C0176A1 (PulA^-^) genome can be found under the following accession number JAEDCG000000000. The metagenomic and metatranscriptomic datasets used in this study can be found under the Bioproject PRJNA797778. All source data that support the findings of this study will be available in a data repository at synapse.org upon publication.

