## Supplementary Information for "Identification and characterization of bacterial glycogen-degrading enzymes in the vaginal microbiome"

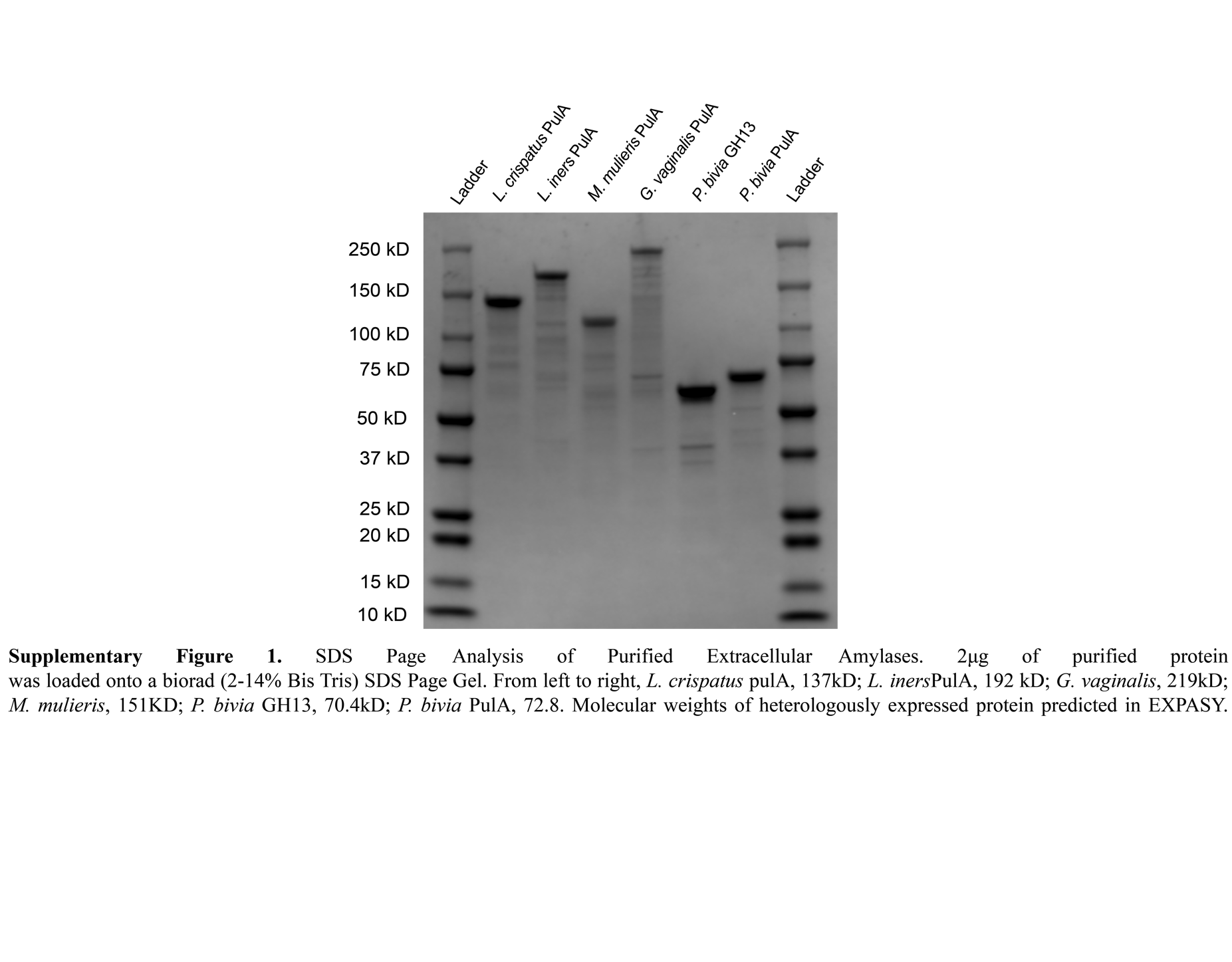

**Supplementary Figure 1.** SDS page analysis of purified extracellular GDEs. 2 μg of purified protein was loaded onto a Biorad (2-14% Bis-Tris) SDS page gel. From left to right, Precision Plus Protein All Blue Standards (BioRad); *L. crispatus* PulA, 137kD; *L. iners* PulA, 192kD; *M. mulieris* PulA, 151kD; *G. vaginalis* PulA, 219kD; *P. bivia* GH 13, 70.4kD; *P. bivia* PulA, 72.8; Precision Plus Protein All Blue Standards (BioRad). Molecular weights of heterologously expressed proteins were predicted in EXPASY.

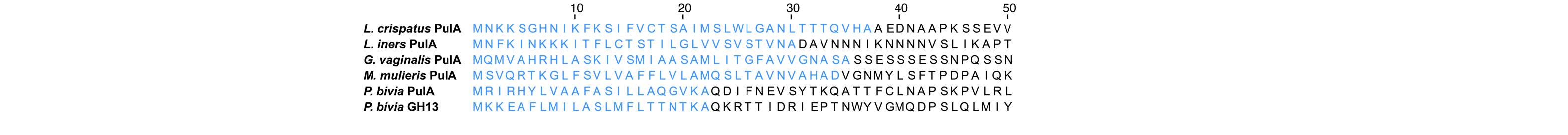

**Supplementary Figure 2.** Signal peptide removal for heterologous expression in *E. coli.* Blue text denotes the amino acid residues removed from each protein sequence during the cloning process.

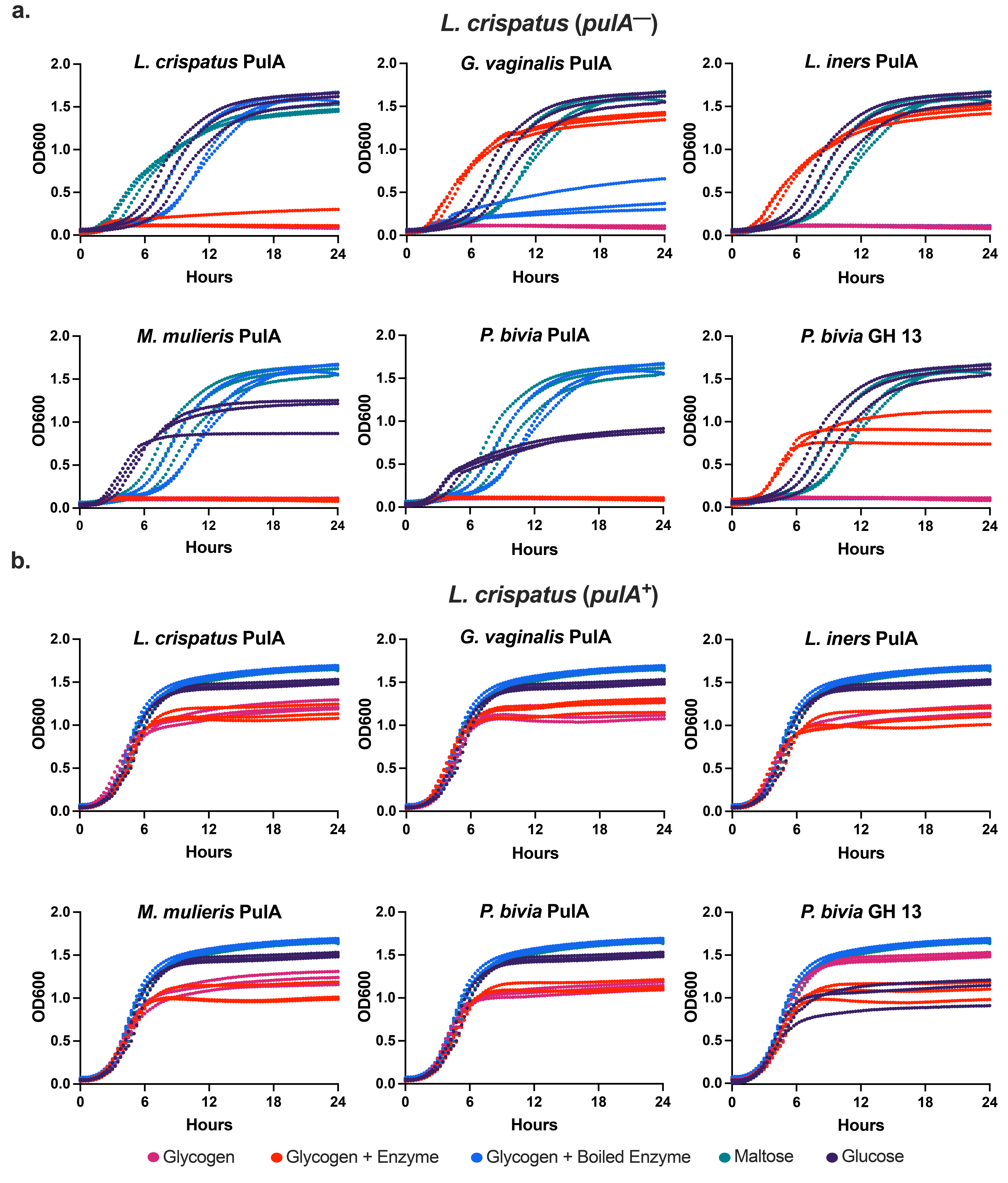
**Supplementary Figure 3.** Recovery of *L. crispatus* growth on glycogen with putative PulA homolog addition. **a.** Growth of *L. crispatus* C0176A1 (*pulA^—^*) containing all relevant controls for protein complementation assays b. Growth of *L. crispatus* MV-1A-US (*pulA^+^*) containing all relevant controls for protein complementation assays. Glycogen, maltose, and glucose conditions are derived from the same data across all graphs within each panel. All growth data consists of three biological replicates performed over three days with three or four technical replicates each day.

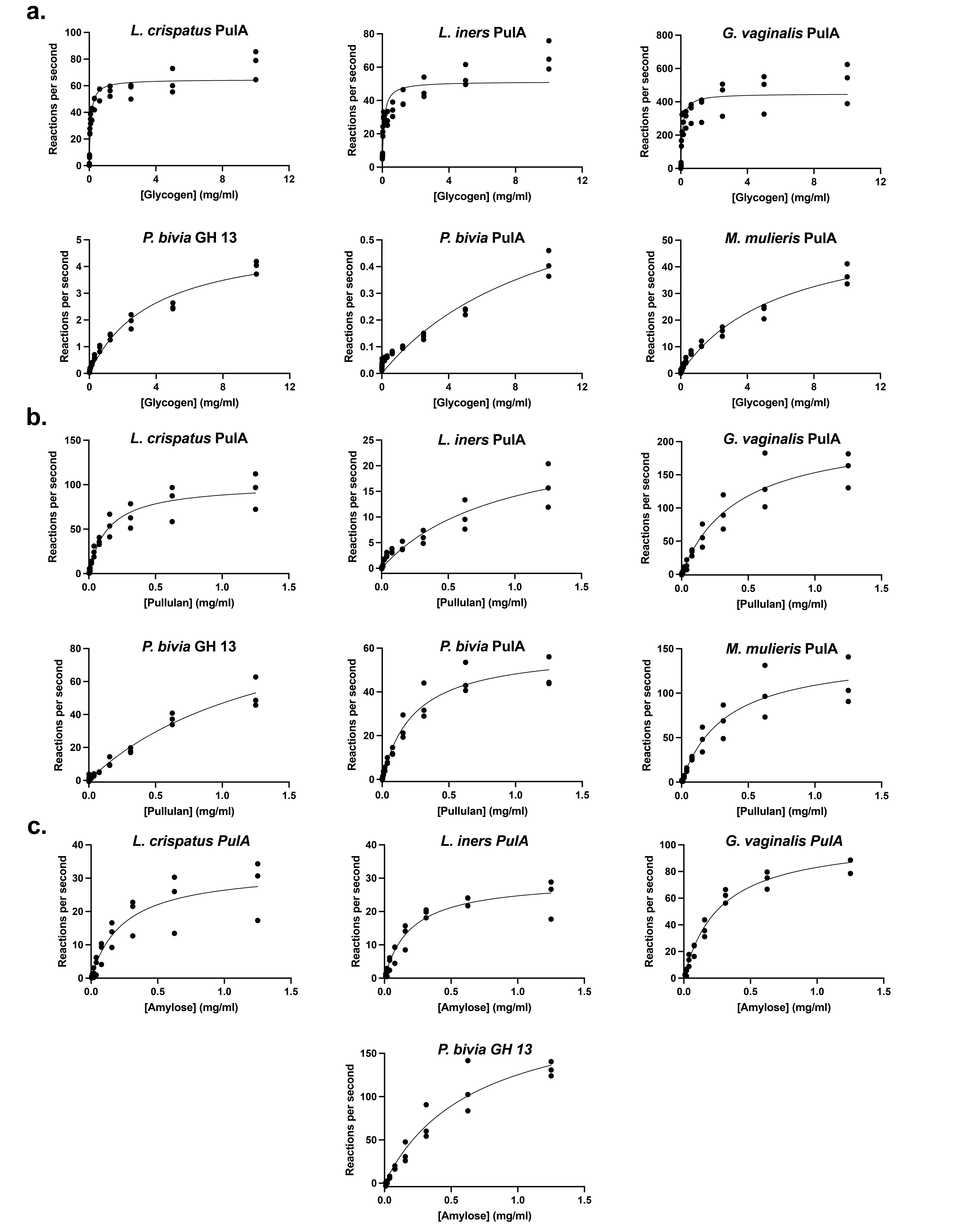
**Supplementary Figure 4.** Kinetic analysis of vaginal microbial GDEs. **a.** Michaelis–Menten kinetic analysis for assays with glycogen. **b.** Michaelis–Menten kinetic analysis for assays with pullulan **c.** Michaelis–Menten kinetic analysis for assays with amylose. All data are representative of three experimental replicates performed over two days. Nonlinear fitting was performed in Graphpad Prism 8.

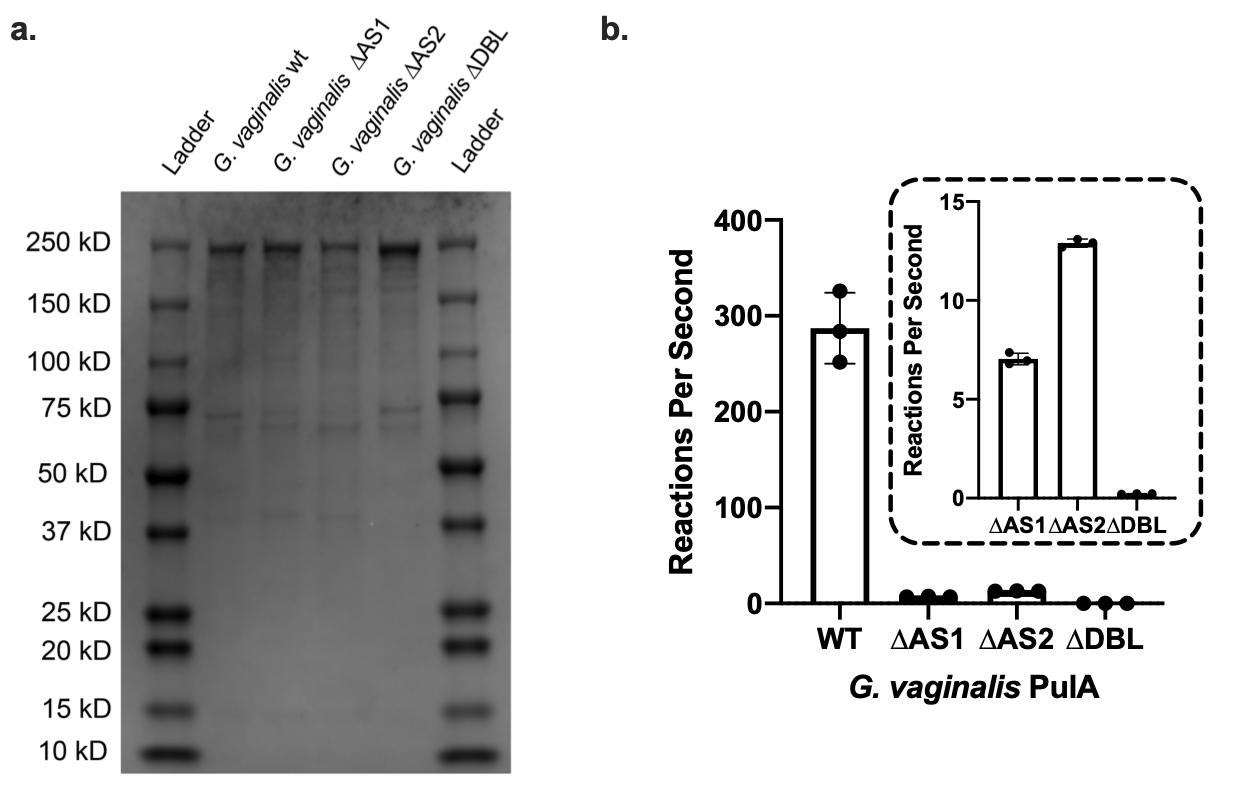

**Supplementary Figure 5.** Active site mutants of *G. vaginalis* PulA. **a.** 2 μg of active site mutants and wild type *G. vaginalis* PulA was loaded onto a Biorad (2-14% Bis-Tris) SDS page gel**. b.** Specific activity of active site mutants on glycogen. Inset graph shows specific activity data for *G. vaginalis* mutants. ∆AS1, ∆AS2, and ∆DBL represents D233A, D1317A, and both mutations respectively. Data is representative of three experimental replicates over two days. Error bars represent one standard deviation above and below the mean.
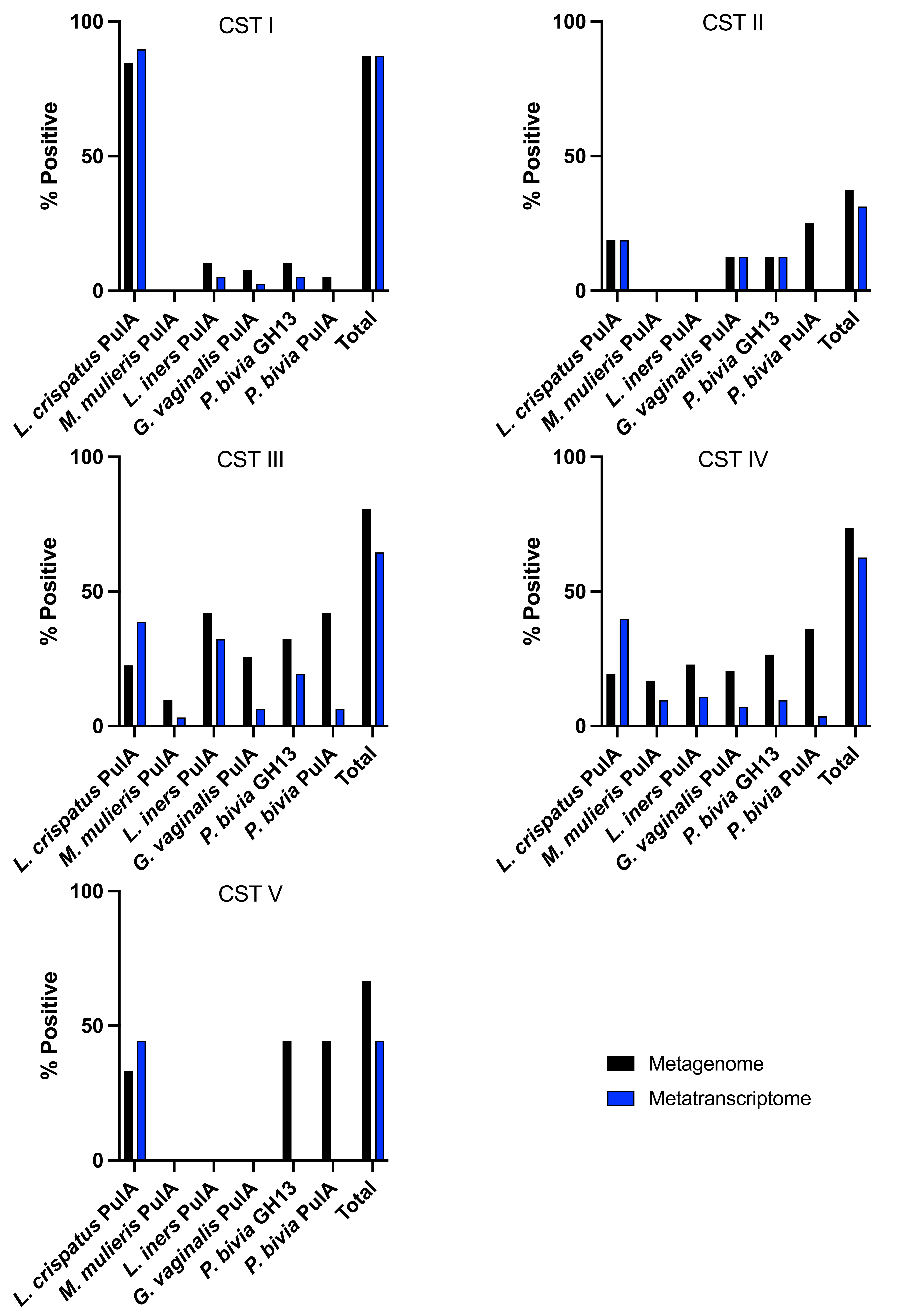
**Supplementary Figure 6.** Microbial GDE metatranscriptomic and metagenomic presence. **% Positive** represents the percentage of samples that contained reads mapping to our query proteins (reads > 0). The sample size is as follows: **CST I**, n = 39; **CST II**, n = 16; **CST III**, n = 31; **CST IV**, n = 83; **CST V**, n = 9.

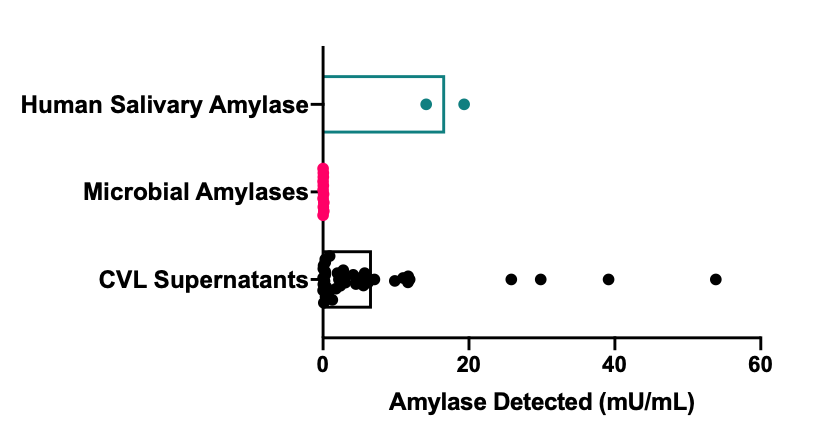

**Supplementary Figure 7.** CVL samples contain human amylase. ELISA detection for human pancreatic amylase in patient CVL supernatants, purified microbial GDEs (1 μM), and purified human salivary amylase (1 μM). 20 CVLs, six different microbial GDEs, and one human salivary amylase were measured in two experimental replicates over two days. All samples and replicates were plotted, and the mean of the samples is denoted by the bar.

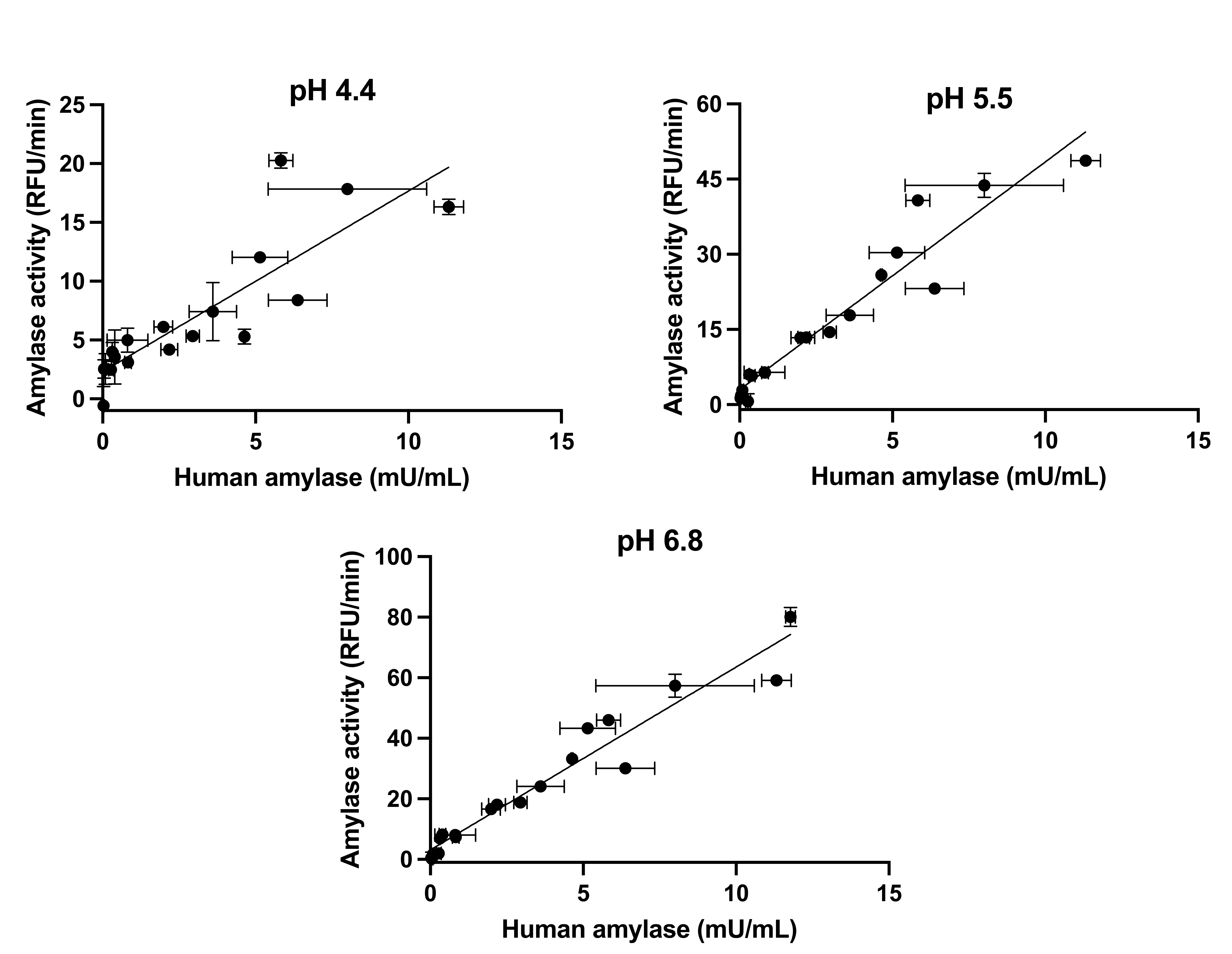

**Supplementary Figure 8.** Activity of human CVL supernatants correlates with human amylase levels detected by ELISA. Outliers were determined and removed using a ROUT test. A Pearson test was used to determine correlation (pH 4.4, r = 0.8556, 95% CI 0.6472 to 0.9450, p = <0.0001; pH 5.5, r = 0.9611, 95% CI 0.8966 to 0.9857, p = <0.0001; pH 6.8, r = 0.9713, 95% CI 0.9252 to 0.9891, p = <0.0001) Activity data represents the mean of three experiments over two days and the ELISA detection is the mean of two experiments over two days. Error bars represent one standard deviation above and below the mean.

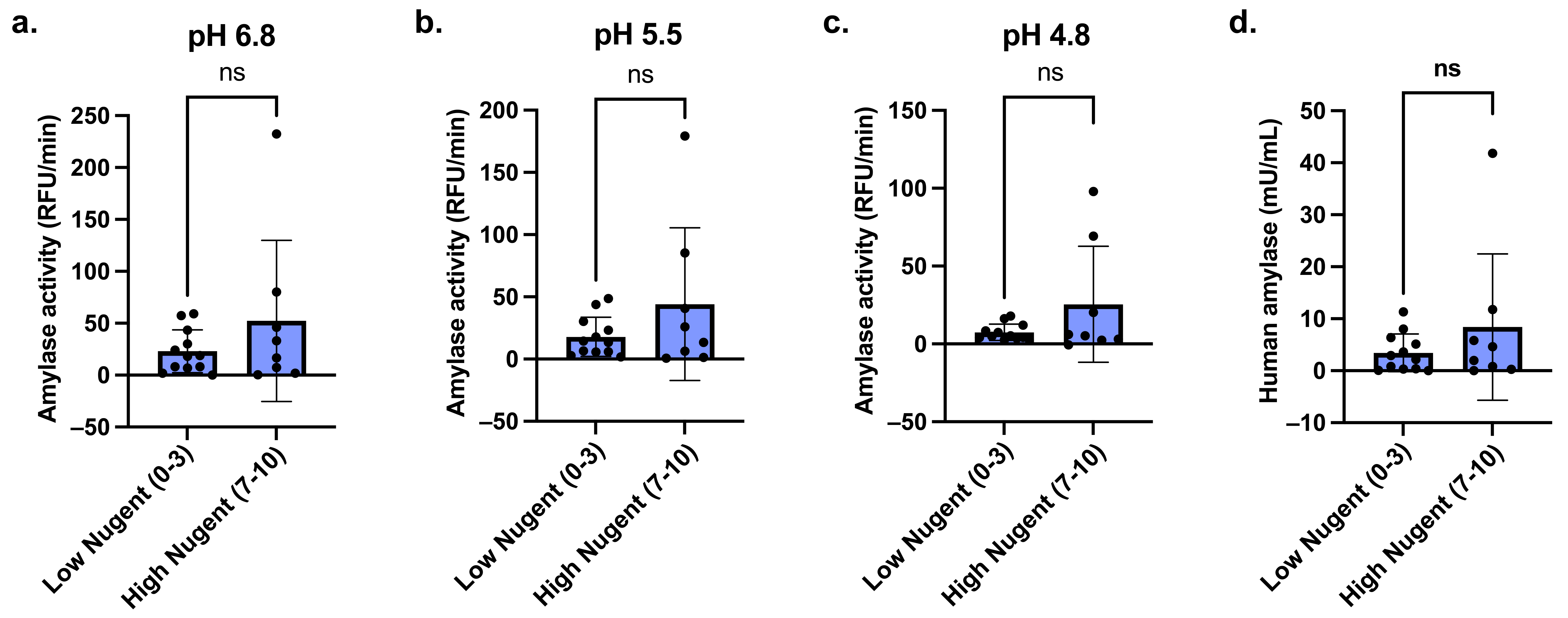

**Supplementary Figure 9.** Human amylase presence and amylase activity do not correlate with Nugent score. **a.** CVL activity levels measured using a fluorescent starch substrate at pH 6.8 stratified by Nugent score determination (P = 0.2249) **b.** CVL activity levels measured using a fluorescent starch substrate at pH 5.5 stratified by Nugent score determination (P = 0.1702) **c.** CVL activity levels measured using on a fluorescent starch substrate at pH 4.8 stratified by Nugent score determination (P = 0.1105) **d.** ELISA amylase detection of human amylase in CVL samples stratified by Nugent score determination (P = 0.2535). The X-axes of all plots are Nugent scores (Low Nugent (0-3), n=12; High Nugent (7-10), n=8). Error bars are representative of one standard deviation above and below the mean. A non-parametric t-test was used to determine statistical significance. P-value symbols: P>0.05 (ns), P≤0.05 (*), P≤0.01 (**), P≤0.001 (***), P≤0.0001(****). All plots were created, and statistics performed in GraphPad Prism 9.

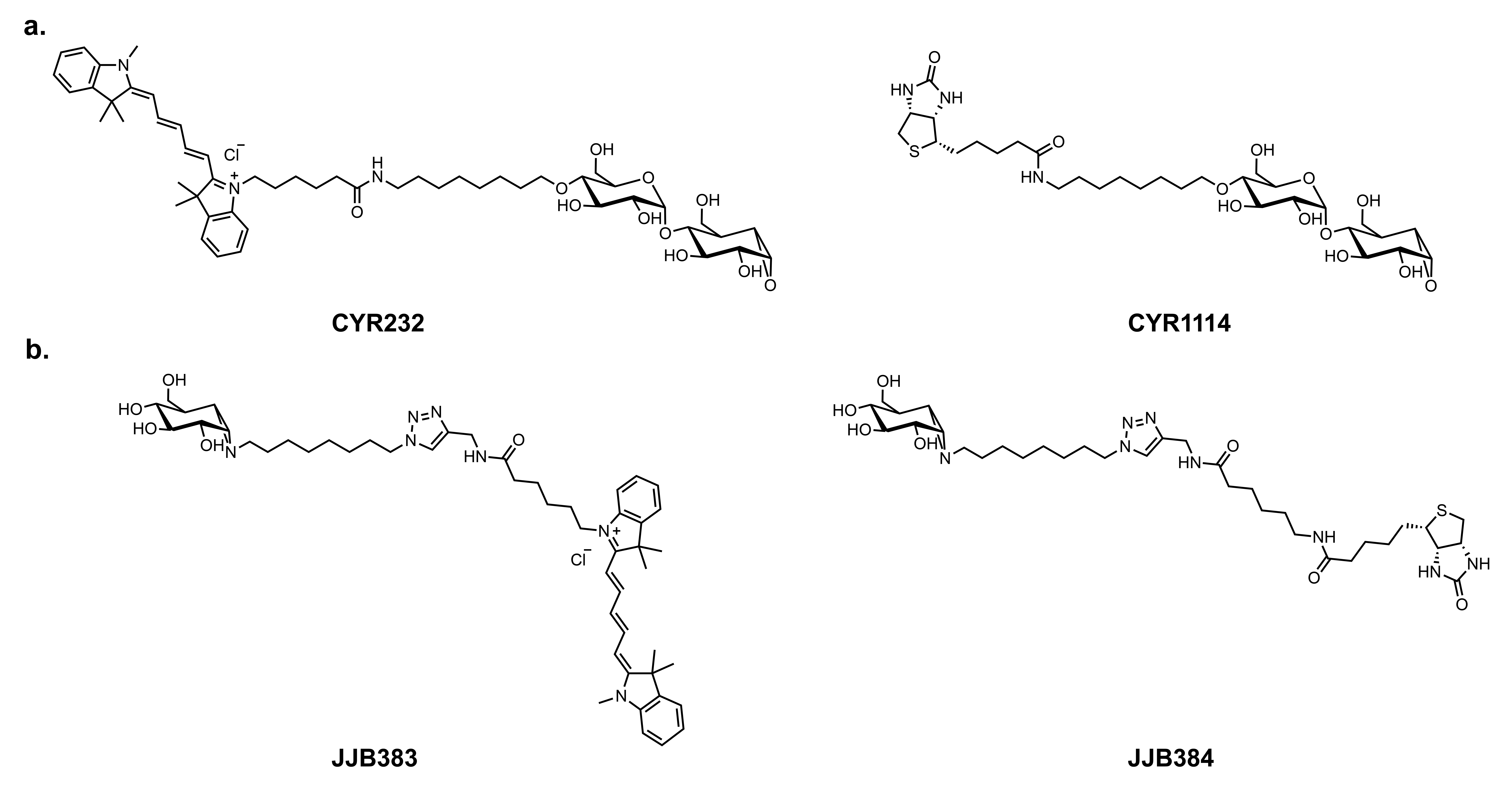

**Supplementary Figure 10.** Chemical structures of the probes used for activity-based protein profiling **a.** α-amylase probes (Amy-ABP) with Cy5 (left) and biotin (right) handles. **b.** α-glucosidase probes (Glc-ABP) with Cy5 (left) and biotin (right) handles.

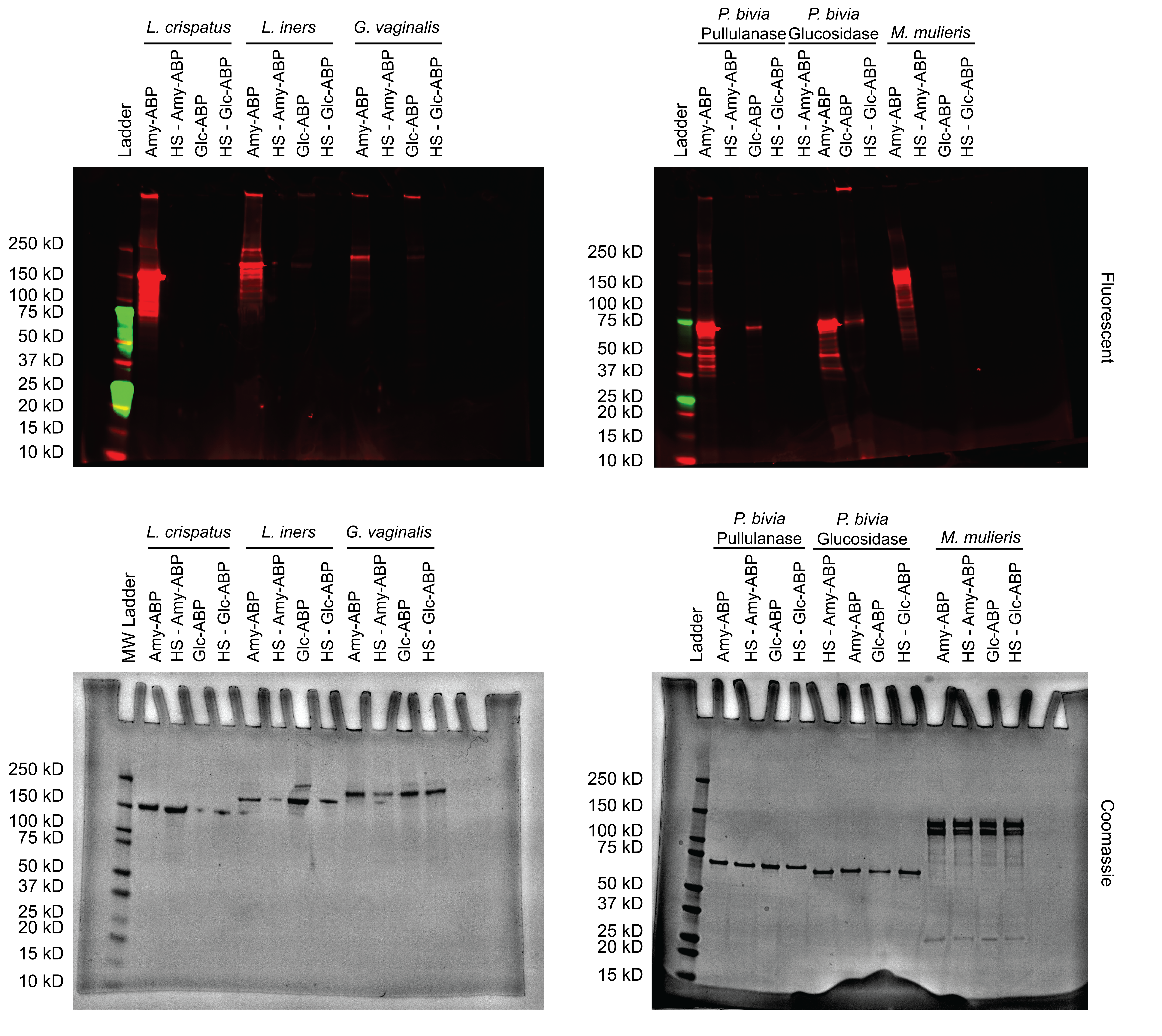

**Supplementary Figure 11.** Purified GDEs from vaginal bacteria react with ABPP probes. Purified GDEs were probed using fluorescent Amy-ABP, Glc-ABP, or vehicle only (**NP**, no probe; **HS**, heat shock) for 2 hours at 37^o^ C and visualized on a 4-20% SDS-PAGE gel (Bio-Rad). Left lane: Precision Plus Protein WesternC standard (Bio-Rad). Fluorescence gel images (top) and Coomassie-stained image of the same gels (bottom) are representative of three independent experiments over three days.

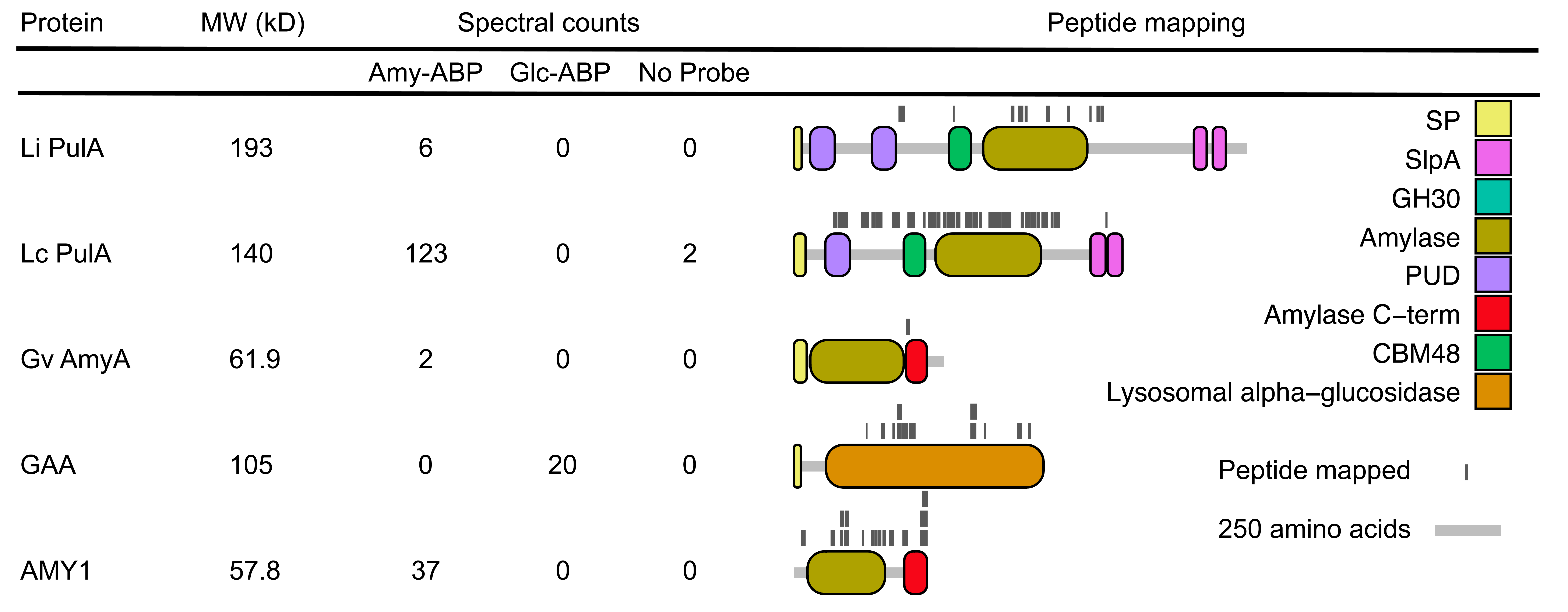

**Supplementary Figure 12.** Peptide mapping to proteins identified through ABPP. Spectral counts indicate the sum of total spectra assignable to a protein across all six individual samples. Abbreviations: **Li PulA**, *L. iners* PulA; **Lc PulA**, *L. crispatus* PulA; **Gv AmyA**, *G. vaginalis* AmyA; **GAA**, human lysosomal α-glucosidase; **AMY1**, human salivary amylase; **SP**, signal peptide; **SlpA,** surface layer protein A; **GH**, glycoside hydrolase domain **PUD,** bacterial pullulanase-associated domain; **CBM,** Carbohydrate binding module.

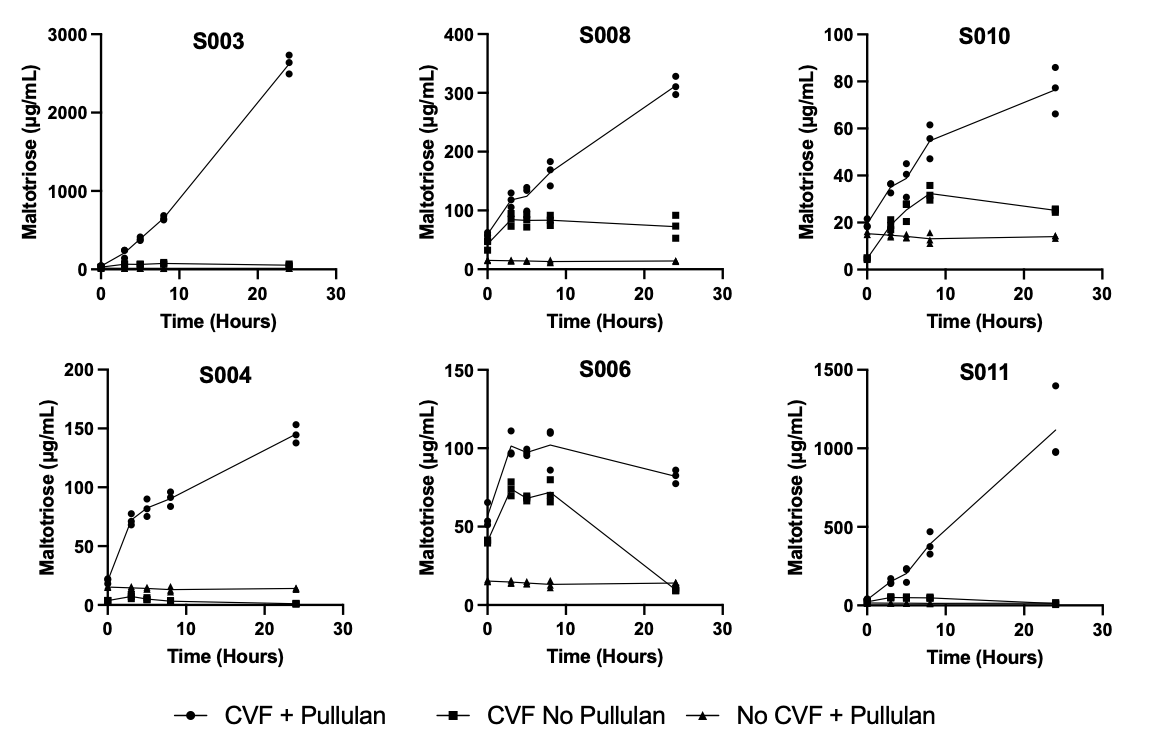

**Supplementary Figure 13.** Time-course analysis of CVF sample pullulan degradation confirms enzymatic activity. Timepoints from CVF reactions were removed and frozen at 3, 5, 8, and 24 hours. These data are derived from three experimental replicates over two days.

**Supplementary Table 1:** Strains and Plasmids Used in this Study

| Name | Description / Use | Reference / Source |
| --- | --- | --- |
| Strains | | |
| E. coli DH5α | Cloning strain | NEB (New England Biolabs) |
| *E. coli* BL21(DE3) | Protein expression and purification | NEB |
| *E. coli* Arctic Express (DE3) | Protein expression with co-expression of psycrophilic chaperones | Agilent |
| *Lactobacillus crispatus* MV-1A-US | Source of *pulA* gene, Growth Studies | BEI |
| Lactobacillus iners LEAF 3008A-a | Source of *pulA* gene, Growth Studies | BEI |
| Gardnerella vaginalis JCP7276 | Source of *pulA* gene, Growth Studies | BEI |
| Mobiluncus mulieris str. 28-1 | Source of *pulA* gene, Growth Studies | BEI |
| Prevotella bivia GED7760C | Source of *pulA* gene, Growth Studies | BEI |
| Lactobacillus crispatus C0176A1 | Growth Studies | VMRC |
| Plasmids | | |
| pET28a | ColE1 lacI pT7 Kn^R^. Expression vector | Novagen |
| pETpullLC | pET28 with N-HIS-tagged *L. crispatus pulA* | This work |
| pETpullGV | pET28 with N-HIS-tagged *G. vaginalis pulA* | This work |
| pETpullLI | pET28 with N-HIS-tagged *L. iners pulA* | This work |
| pETpullMM | pET28 with N-HIS-tagged *M. mulieris pulA* | This work |
| pETpullPB | pET28 with N-HIS-tagged *P. bivia pulA* | This work |
| pETgh13PB | pET28 with N-HIS-tagged *P. bivia glycosidase* | This work |
| pETpullGV-AS1 | pETpullGV with D233A mutation (based on numbering of native protein sequence) | This work |
| pETpullGV-AS2 | pETpullGV with D1317A mutation | This work |
| pETpullGV-DM | pETpullGV with D233A and D1317A mutations | This work |

**Supplementary Table 2:** Cloning Primers Used in this Study

| Fragment | Template | Primer | Sequence |
| --- | --- | --- | --- |
| pETpullLC | | | |
| *L. crispatus* *pulA* | gDNA | oBMW_005  oBMW_006 | gcctggtgccgcgcggcagcGCAGAGGACAATGCAGCTC  cagcttcctttcgggctttgCTAACCAATTCTTTTAACATTTCTAGCC |
| Backbone | pET28 | oBMW_003  oBMW_004 | GCTGCCGCGCGGCACCAG  CAAAGCCCGAAAGGAAGCTG |
| pETpullGV | | | |
| *G. vaginalis* *pulA* | gDNA | oBMW_007  oBMW_008 | cagcggcctggtgccgcgcggcagcAGTAGTGAGTCTAGTAGTG  caactcagcttcctttcgggctttgCTATCTAAATGCAACTTTATAACG |
| Backbone | pET28 | oBMW_003  oBMW_004 | GCTGCCGCGCGGCACCAG  CAAAGCCCGAAAGGAAGCTG |
| pETpullLI | | | |
| *L. iners* *pulA* | gDNA | oBMW_009  oBMW_062 | ctttaagaaggagatatacc**ATG**ACCCACCTGAACATCGC  cctcgtttcttggttcttcgTTACATCGCCGCAGCGAAGATTG |
| Backbone | pET28 | oBMW_003  oBMW_004 | GCTGCCGCGCGGCACCAG  CAAAGCCCGAAAGGAAGCTG |
| pETpullPB | | | |
| *P. bivia* *pulA* | gDNA | oBMW_168  oBMW_169 | gcctggtgccgcgcggcagcCAGGATATTTTCAACGAAGTG  cagcttcctttcgggctttgTTAGTTGTGTAAGATCAGTG |
| Backbone | pET28 | oBMW_003  oBMW_004 | GCTGCCGCGCGGCACCAG  CAAAGCCCGAAAGGAAGCTG |
| pETgh13PB | | | |
| *P. bivia* *glycosidase* | gDNA | oBMW_170  oBMW_171 | gcctggtgccgcgcggcagcCAAAAAAGAACTACTATCGATC  cagcttcctttcgggctttgTTAGAACTCTAATATCATCACAC |
| Backbone | pET28 | oBMW_003  oBMW_004 | GCTGCCGCGCGGCACCAG  CAAAGCCCGAAAGGAAGCTG |
| pETpullMM | | | |
| *M. mulieris* *pulA* | gDNA | oBMW_184  oBMW_185 | gcctggtgccgcgcggcagcGATGTTGGGAATATGTACC  cagcttcctttcgggctttgCTACTTGGTAAACATTGGTAG |
| Backbone | pET28 | oBMW_003  oBMW_004 | GCTGCCGCGCGGCACCAG  CAAAGCCCGAAAGGAAGCTG |
| pETpullGV-AS1 | | | |
| *G. vaginalis pulA* Fragment A | pETpullGV | oBMW_209  oBMW_210 | atcatcacagcagcggcctggtgccgcgcggcagcAGTAGTGAGTCTAGTAGTGAATC  ggagaaatatgttttactgcagcGATACGGAAGCCAGCAAC |
| *G. vaginalis* *pulA* Fragment BC | pETpullGV | oBMW_211  oBMW_214 | ttgctggcttccgtatcgctGCAGTAAAACATATTTCTCCAAC tggcagcagccaactcagcttcctttcgggctttgCTATCTAAATGCAACTTTATAACG |
| Backbone | pET28 | oBMW_003  oBMW_004 | GCTGCCGCGCGGCACCAG  CAAAGCCCGAAAGGAAGCTG |
| pETpullGV-AS2 | | | |
| *G. vaginalis pulA* Fragment AB | pETpullGV | oBMW_209  oBMW_212 | atcatcacagcagcggcctggtgccgcgcggcagcAGTAGTGAGTCTAGTAGTGAATC  tcaatcaatcccatcaaagcAAAACGGAAGCCGTCAATAC |
| *G. vaginalis pulA* Fragment C | pETpullGV | oBMW_213  oBMW_214 | gtattgacggcttccgttttgctTTGATGGGATTGATTGAC  tggcagcagccaactcagcttcctttcgggctttgCTATCTAAATGCAACTTTATAACG |
| Backbone | pET28 | oBMW_003  oBMW_004 | GCTGCCGCGCGGCACCAG  CAAAGCCCGAAAGGAAGCTG |
| pETpullGV-DM | | | |
| *G. vaginalis pulA* Fragment A | pETpullGV | oBMW_209  oBMW_210 | atcatcacagcagcggcctggtgccgcgcggcagcAGTAGTGAGTCTAGTAGTGAATC  ggagaaatatgttttactgcagcGATACGGAAGCCAGCAAC |
| *G. vaginalis pulA* Fragment B | pETpullGV | oBMW_211  oBMW_212 | ttgctggcttccgtatcgctGCAGTAAAACATATTTCTCCAAC  tcaatcaatcccatcaaagcAAAACGGAAGCCGTCAATAC |
| *G. vaginalis pulA* Fragment C | pETpullGV | oBMW_213  oBMW_214 | gtattgacggcttccgttttgctTTGATGGGATTGATTGAC  tggcagcagccaactcagcttcctttcgggctttgCTATCTAAATGCAACTTTATAACG |
| Backbone | pET28 | oBMW_003  oBMW_004 | GCTGCCGCGCGGCACCAG  CAAAGCCCGAAAGGAAGCTG |

**Supplementary Table 3:** LC-MS compound specific detection parameters

| **Compound** | **Mass *(m/z)*** | **Cone Voltage (volts)** | **Elution time (min)** |
| --- | --- | --- | --- |
| Glucose | 203.0 | 47 | 3.53 |
| Maltose | 365.0 | 76 | 3.85 |
| Maltotriose | 527.1 | 98 | 4.03 |
| Maltotetraose | 689.1 | 100 | 4.15 |
| Maltopentaose | 851.3 | 100 | 4.28 |

**Supplementary Table 4:** Metadata describing CVL sample donors used in human amylase ELISA and amylase activity assays.

| **Age** | **Race** | **Ethnicity** | **Diagnosis** | **Nugent Score** | **Contraception** |
| --- | --- | --- | --- | --- | --- |
| 27 | White | Non-Hispanic | Normal exam | 0 | None/Rhythm |
| 30 | White | Non-Hispanic | Vulvodynia | 3 | None/Rhythm |
| 27 | White | Non-Hispanic | BV | 7 |  |
| 25 | White | Non-Hispanic | Vulvar dermatitis | 1 | OCP |
| 25 | White | Non-Hispanic | BV | 8 | Nexplanon |
| 46 | White | Non-Hispanic | BV | 7 | None/Rhythm |
| 39 | White | Hispanic | Vulvodynia | 1 | None/Rhythm |
| 49 | White | Non-Hispanic | BV | 8 | None/Rhythm |
| 22 | White | Non-Hispanic | Idiopathic sx | 0 |  |
| 37 | White | Non-Hispanic | Normal exam | 0 | None/Rhythm |
| 25 | White | Non-Hispanic | BV | 8 | None/Rhythm |
| 29 | White | Non-Hispanic | BV | 7 | None/Rhythm |
| 46 | White | Non-Hispanic | BV | 7 | Sterilization |
| 63 | White | Non-Hispanic | Menopausal symptoms | 8 |  |
| 25 | White | Non-Hispanic | Vulvar dermatitis | 0 | Mirena/Skyla |
| 21 | White | Non-Hispanic | Normal exam | 0 | OCP |
| 45 | White | Non-Hispanic | Normal exam | 2 |  |
| 24 | White | Non-Hispanic | Yeast | 0 | OCP |
| 27 | White | Non-Hispanic | Yeast | 0 |  |
| 36 | White | Non-Hispanic | Yeast | 2 | None/Rhythm |

**Supplementary Table 5:** Metadata describing CVF sample donors used in ABPP and pullulanase assays.

| **Sample** | **Age Range** | **Race/Ethnicity** | **Hormonal Contraceptive Use** | **Day of Cycle** |
| --- | --- | --- | --- | --- |
| S003 | 18-25 | Hispanic/White | No | 9 |
| S004 | 18-25 | White | No | 16 |
| S006 | 18-25 | White | No | 21 |
| S008 | 18-25 | White | Yes | 15 |
| S010 | 18-25 | Asian/White | Yes | NA |
| S011 | 18-25 | Hispanic | Yes | NA |
